# Distributed sensory coding by cerebellar complex spikes in units of cortical segments

**DOI:** 10.1101/2020.09.18.301564

**Authors:** Takayuki Michikawa, Takamasa Yoshida, Satoshi Kuroki, Takahiro Ishikawa, Shinji Kakei, Shigeyoshi Itohara, Atsushi Miyawaki

## Abstract

Sensory processing is essential for motor control. Climbing fibers from the inferior olive transmit sensory signals to Purkinje cells, but how the signals are represented in the cerebellar cortex remains elusive. We examined the olivocerebellar organization of the mouse brain by optically measuring complex spikes (CSs) evoked by climbing fiber inputs over the entire dorsal surface of the cerebellum. We discovered that the surface was divided into approximately 200 segments each composed of ∼100 Purkinje cells that fired CSs synchronously. Our *in vivo* imaging of evoked responses revealed that whereas stimulation of four limb muscles individually similar global CS responses across nearly all segments, the timing and location of a stimulus were derived by Bayesian inference from coordinated activation and inactivation of multiple segments on a single trial basis. Our findings suggest that the cerebellum performs segment-based distributed population coding by assembling probabilistic sensory representation in individual segments.

## INTRODUCTION

The cerebellum plays a pivotal role in motor control (Ito, 1984) and perceptual processing (Baumann et al., 2015). For the execution of these tasks, it receives a great deal of information about the external world and the state of the body on a moment-to-moment basis (Rondi-Reig et al., 2014). Sensory information is considered to be essential not only for controlling ongoing movements but also for updating internal models of the external world and the motor apparatus using sensory prediction error-based learning (Shadmehr et al., 2010; Wolpert et al., 1998). However, how sensory information is represented in the cerebellum is less well understood.

Purkinje cells (PCs) are the sole output neurons in the cerebellar cortex that receives two types of excitatory inputs: climbing fibers and mossy fibers relayed by granule cells (Figure S1A). Climbing fibers arise exclusively from the inferior olive (IO) (Figure S1B), which is anatomically divided into several subdivisions. The topographical distribution of climbing (olivocerebellar) fibers in the contralateral cortex from each of the IO subdivisions was characterized as a parasagittally elongated “zone” (Voogd et al., 2013) (Figure S1C). The single input of a climbing fiber induces a complex spike (CS), a short burst of impulses, in a postsynaptic PC. On the other hand, a PC receives inputs from granule cells and exhibit changes in the firing rate of simple spikes (Figure S1A).

Both climbing and mossy fibers transmit sensory signals from peripheral tissues, but somatotopy, the point-to-point correspondence of an area of the body to a specific point on the central nervous system, has not yet been well established in the cerebellar cortex (Manni and Petrosini, 2004). The cerebellar homuncular somatotopy that is illustrated in common textbooks (Purves et al., 2018; Purves et al., 2007) represents separate bodily parts in the anterior lobe and paramedian lobules of the cerebellum (Figure S1D). It was originally proposed based on evoked potentials recorded from the surface of the cerebellar cortex in barbiturate- or chloralosan-anesthetized cats (Snider and Stowell, 1944). To detect evoked potentials on the cerebellar surface efficiently under unanesthetized state, in contrast, a subsequent study used decerebrated cats but observed no somatotopic organization in the anterior lobe of the cerebellum (Combs, 1954) (Figure S1E, left); discrete localization of potentials was only observed after the injection of sodium pentobarbital in the study (Figure S1E, right). Based on the long latency, these surface potentials were later interpreted as climbing fiber responses (Eccles et al., 1967), suggesting that the aforementioned studies examined the somatotopic termination of spino-olivocerebellar pathways. The somatotopy in the cat cerebellum was further investigated using differently operated preparations (Oscarsson, 1968); the surface potential signals of each spino-olivocerebellar pathway were enhanced by transecting the unrelated spinal cord regions. This study localized evoked signals in accordance with the parasagittal zones (Figure S1C) to propose a new type of somatotopy “longitudinal somatotopy” in the anterior lobe (Figure S1F). With the times, however, the importance of intact neuraxis has been underscored for somatotopy assessment. Without any surgery or anesthetization on cats, a later study that used single-unit recordings for observing climbing fiber responses found no somatotopic organization in the cerebellar anterior lobe (Leicht and Schmidt, 1977) (Figure S1G). Taken together, the existence of the somatotopic organization of spino-olivocerebellar pathways has long been controversial, even if the discussion is limited to the anterior lobe of the cat cerebellum where the sensory homunculus is believed to exist.

Investigation of elementary units in the cerebellum is critical for understanding modular structure of the cerebellar circuits. The theory of modularity (or localization) argues that the brain is organized into discrete modules, each of which processes a particular type of information. By contrast, the theory of distributed processing argues that any information is processed by multiple regions of the brain, and that any brain region represents multiple types of information. Although single-unit recordings can be used to examine with single-cell resolution how sensory signals are converged or diverged in the brain, this approach usually measures only a limited number of neurons in one experiment. Local examination of spino-olivocerebellar inputs revealed several “microzones” in a parasagittal zone of the cat cerebellum (Figure S1H) (Andersson and Oscarsson, 1978). Microzones were found to receive a combination of various peripheral inputs and define components of olivo-cortico-nuclear microcomplexes, which have been proposed as the functional units in the cerebellum (Apps and Garwicz, 2005; Ito, 1984; Oscarsson, 1980). However, the observed areas were so small that the modularity was discussed just inside a zone. It was also found that PCs showing similar receptive fields on the body of intact cats were clustered within a lobule to make “patches” or “mosaic” pattern (Figure S1I) (Eccles, et al., 1968; Leicht and Schmidt 1977; Robertson et al., 1982). However, the lobule regions were less densely populated by recorded PCs, and it was difficult to clearly delineate individual functional units. On the other hand, cerebellar modularity may be related to the heterogeneity of PCs in their biochemical compositions. “Stripes” or “bands” have been defined throughout the cerebellar cortex based on gene expression patterns of some molecular markers of PC populations, such as phospholipase C b4 and aldolase C (Figure S1J) (Apps and Hawkes, 2009). However, the functional significance of these histological compartmentalizations is not clear.

Within the central nervous system, a climbing fiber shows a unique characteristic; its firing occurs at an extremely low rate (∼1 Hz). Because at most one, if any, CS can be detected from single PCs during natural movements, such as reaching (Keating and Thach, 1995; Kitazawa et al., 1998) and eye movements (Catz et al., 2005), spike organization cannot be analyzed adequately according to the traditional rate coding scheme, in which a substantial amount of information is extractable from the changing firing rate of single neurons. In fact, it has been debated for several decades what kind of information is encoded by such sparsely firing climbing fibers (Lang et al., 2017).

At all events, comprehensive understanding of whether representation of each body part is localized or distributed in the cerebellar cortex awaits a new method for wide mapping of CSs with high spatiotemporal resolution. In this study, we established a method for real-time *in vivo* optical measurements of CS activity from more than 10,000 PCs simultaneously, and then applied probability theory to decipher large-scale CS population codes in single-trial experiments. Our study has unveiled a distributed moment-by-moment way in which the olivocerebellar system represents the conditional probability of the occurrence of every sensory event in a large population of CSs through its novel hierarchical organizations, leading to a drastic change in our understanding about the operational principle of the cerebellum.

## RESULTS

### Discovery of olivocerebellar segments

The generation of a CS results in a massive increase in Ca^2+^ in PC dendrites. Aiming for reliable detection of CSs, we adopted the ratiometric measurement of Ca^2+^; we have been employing yellow cameleon 2.60 (YC2.60), a genetically-encoded Ca^2+^ sensor based on fluorescent resonance energy transfer (FRET) between cyan- and yellow-emitting fluorescent proteins (Miyawaki et al., 1997; Nagai et al., 2004). We previously established an adenoviral transduction method for targeting YC2.60 expression to PCs of the mouse cerebellum (Horikawa et al., 2010; Yamada et al., 2011). Because the targeting efficiency was 30–50%, YC2.60-expressing PCs were well delineated individually. Using living mice, we first performed simultaneous measurements of dendritic signals of YC2.60 and extracellular potential fields to confirm that every increase in the FRET signals exclusively reflected occurrence of a CS (Figures S2A–S2D). To monitor CSs in all PCs that were optically accessible, we next generated L7-YC2.60 mice by crossing tLSL-YC2.60 mice (Figure S2E) (Kuroki et al., 2018) with L7-Cre mice; YC2.60 was highly expressed in every PC in the resultant mice (Figure S2F). Additionally, we have developed a special operation to make a large cranial window that covers the entire dorsal surface of the cerebellar cortex (Figure S2G).

We used a custom-made macroscope (Kuroki et al., 2018) (Figure S2H) to monitor YC2.60 signals densely distributed over the cortical surface (Figures 1A and 1B) of an L7-YC2.60 mouse (#1032) (Table S1). Careful registration of cyan and yellow fluorescence images and elaborate processing for generation of their ratio images (see Methods) allowed comprehensive real-time visualization of CS activities of over 10,000 PCs (Figure 1C). Our large-scale measurements uncovered a contiguous transition of ever-changing patterns of both local and global CS synchrony over the cortex. To analyze these complex patterns, we used principal component analysis and spatiotemporal independent component analysis to isolate PC clusters that exhibit synchronous spontaneous CS activity, as reported for the cerebral cortex (Makino et al., 2017). While a programming code was developed previously to isolate single cells within a small field-of-view of two-photon Ca^2+^ imaging data (Mukamel et al., 2009), we modified the code to extend the processing to the entire imaged cortex. Remarkably, this analysis allowed us to divide the cortical surface into 196 segments (Figure 1D), in each of which PCs fired CSs synchronously. Segments in lobule V, simple lobule and crus II exhibited longitudinal shapes, whereas segments in other lobules exhibited irregular shapes located in the rostral or caudal side of the lobule (Figure S2I). This hierarchical structure, consisting of ∼200 CS-associated segments, was observed also in three other L7-YC2.60 mice (Figures S3A to S3C) and was thus hypothesized to reflect functional topographic units of the olivocerebellar system; hereafter these units are referred to as “olivocerebellar segments”.

**Figure 1.**
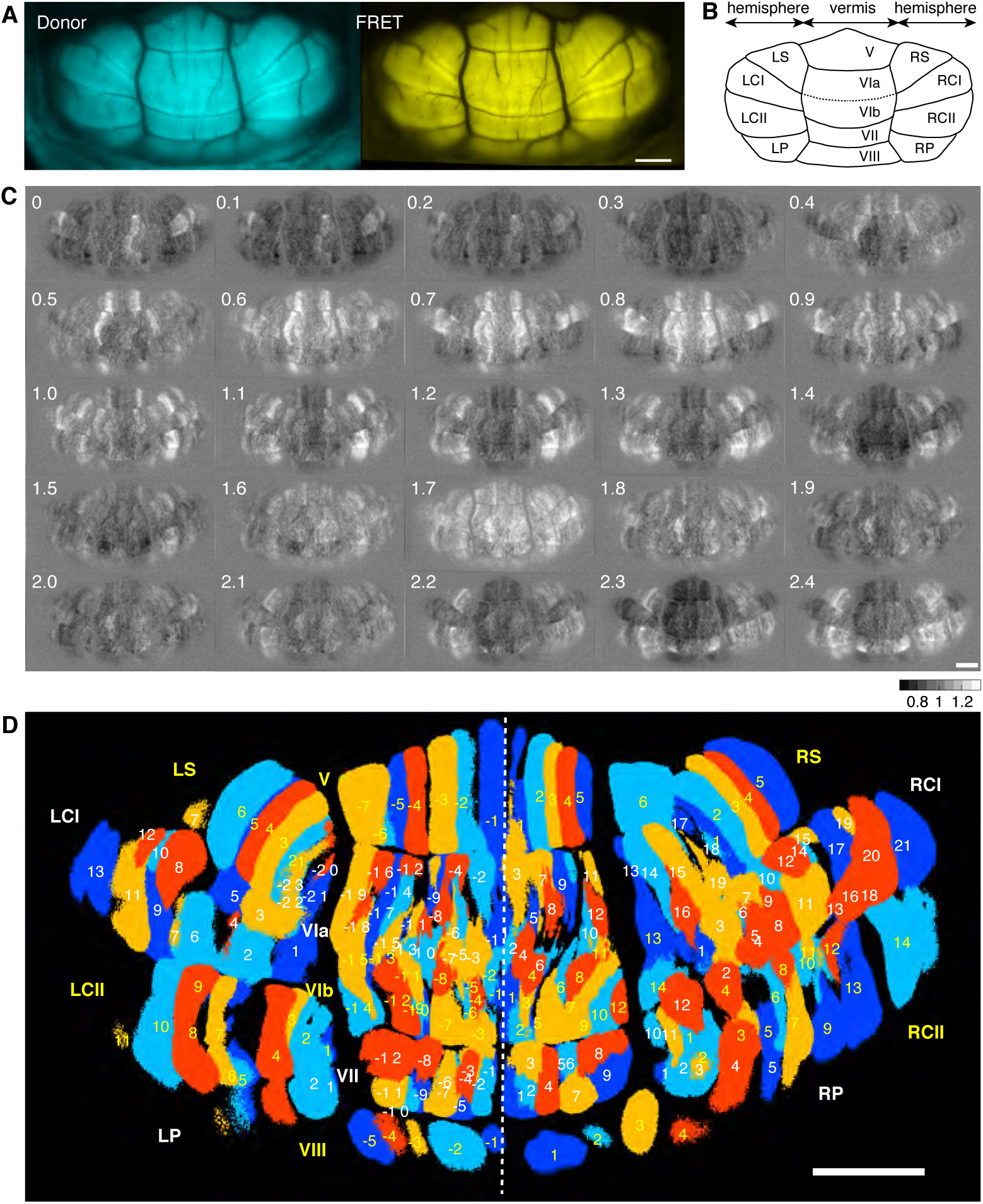
Olivocerebellar segments, functional units of the mouse cerebellar cortex revealed by comprehensive observation of CS activity. (A) Donor and FRET fluorescence images acquired from the dorsal surface of the cerebellum acquired using a custom-made macroscope through a large cranial window. (B) Dorsal view of the mouse cerebellum with macroscopic division into lobules. The following abbreviations are used to label lobules. LCI and RCI: crus I in the left and right hemispheres, respectively. LS and RS: simple lobules in the left and right hemispheres, respectively. LCII and RCII: crus II in the left and right hemispheres, respectively. LP and RP: paramedian lobules in the left and right hemispheres, respectively. (C) Sequential images of mean-subtracted ratio (32-bit floating-point values, see Methods), capturing the spatiotemporal pattern of spontaneous CS activity. Relative time is shown in sec in each image. Background ratio equals 1 (see equation [1] in Methods). (D) A total of 196 olivocerebellar segments identified on the basis of CS synchrony measured by independent component analysis. They are colored in blue, light blue, orange, and red for visual discrimination. Segments in each lobule are numbered in yellow or white according to their cortical location relative to the midline (broken white line). Note that segments in V, LS, and RS extend the entire rostrocaudal length of the lobules. (A, C, and D) Scale bars: 1 mm. Image data are from L7-YC2.60 mouse #1032 (Table S1). See also Figures S2 and S3.

### CS activity over olivocerebellar cross

Temporal profiles of spontaneous CS activities in olivocerebellar segments were comparatively examined throughout the cortex of mouse #1032 (Figure 2A). We first noticed that all 196 segments oscillated at approximately 1 Hz, and analyzed the cross-correlation among their time series (Figure 2B). As a large fraction of the matrix was colored in red or blue for positive or negative cross correlation, respectively, those segments could be categorized into two groups with mutually opposite phases; the first group (labeled in orange) covered the segments located in lobules V–VIII and crus I, whereas the second group (labeled in cyan) covered those in simple lobules, crus II and paramedian lobules. In accordance with this finding, our phase analysis using the Hilbert transform (Figure 2C) clearly showed an in-phase relationship between any two segments inside the first group (VIa-1 vs. VIb-1) or second group (LP3 vs. LS7), but an anti-phase relationship between the two groups (VIa-1 vs. LP3). The whole domain composed of segments of the first group, which spans the vermal and bilateral crus I lobules, forms a cruciform shape over the cortex (Figure 2D). This characteristic structure was identified also in the three other L7-YC2.60 mice (Figures S3D and S3E), and is hereafter referred to as the “olivocerebellar cross”. Two-dimensional heatmaps of inter-segmental cross-correlation coefficients (Figure 2E) illustrate that the phase synchronization was extended over the cortex symmetrically, and that oscillation phase was reversed sharply across the border of the olivocerebellar cross. Of particular interest was the completeness of the remote synchronization at the four corners (outside the cross). On the other hand, global synchronization inside the cross apparently decreased with inter-segmental distance. Such synchrony of CS occurrence was examined locally in previous studies; electrophysiological recording (Llinás and Sasaki, 1989) or two-photon Ca^2+^ imaging (Mukamel et al., 2009) showed that the synchrony decreased with the mediolateral separation distance between PCs sampled within a lobule, such as rodent crus IIa (Llinás and Sasaki, 1989), and mouse lobules V and VI (Mukamel et al., 2009). Since the cross-correlation coefficient in our study reflects the degree of the CS synchrony, we were able to demonstrate such intra-lobular, distance-dependent decay of synchronization more quantitatively and extensively (Figure S4A and Table S2).

**Figure 2.**
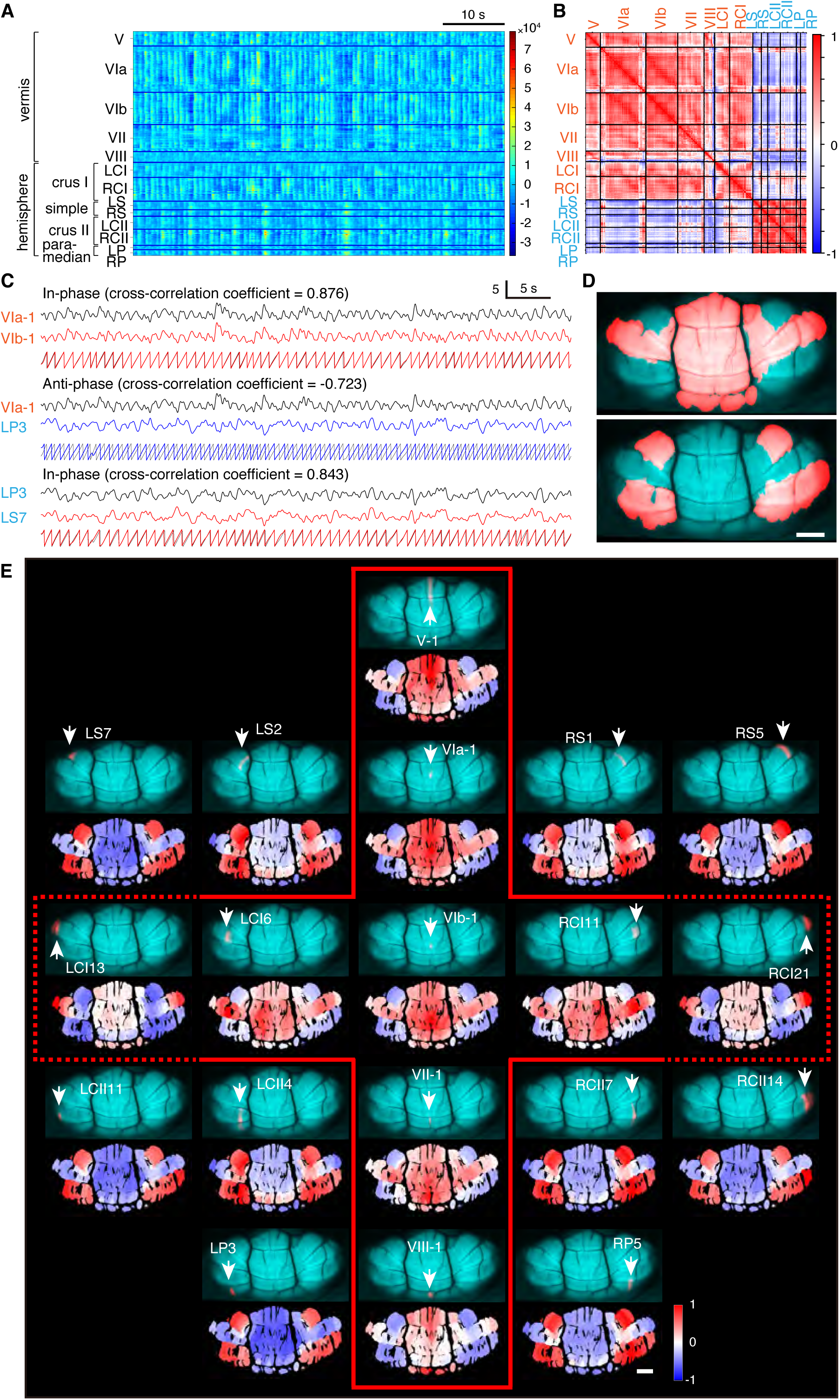
Olivocerebellar cross delineated by phase reversal of spontaneous CS activity of olivocerebellar segments. (A) Temporal profiles of color-coded FRET signals (mean-subtracted ratio, see Methods) from the 196 olivocerebellar segments shown in Figure 1D. Segments in each lobule are aligned according to the mediolateral location of their centroids from left to right. (B) Cross-correlation coefficients (time lag = 0) between segmental FRET signals. Calculated from the data in (A) after normalization (see Methods). Segments/lobules are aligned as in (A). For sharp contrast, vermal lobules and crus I lobules in bilateral hemispheres are combined (orange); the remaining hemispheric lobules are indicated in cyan. (C) Time courses of normalized FRET signals from two segments showing in-phase (VIa-1 vs. VIb-1 and LP3 vs. LS7) or anti-phase (VIa-1 vs. LP3) oscillations. Traces were bandpass-filtered (1.2 ± 0.3 Hz) and processed by the Hilbert transform to generate phase angle (from –π to +π radian) traces (bottom). (D) Segments inside (top) and outside (bottom) the olivocerebellar cross are highlighted in orange against a cyan background. (E) Inter-segmental cross-correlation of normalized FRET signals over the cortex. Nine and ten segments were chosen as references inside and outside the olivocerebellar cross, respectively, and are separated by red lines. Their locations (upper) and heatmaps of cross-correlation coefficients with all segments (lower) are illustrated. Segments/lobules are labeled as in Figure 1D. (B and E) Positive and negative correlations are indicated in red and blue, respectively. (D and E) Scale bars: 1 mm. (A–E) All data are from L7-YC2.60 mouse #1032 (Table S1). See also Figures S3 and S4.

We next analyzed the spatiotemporal pattern of spontaneous CS activities by focusing on their phase at the level of individual segments (Figure S4B). The phase transition dynamics was almost symmetric over the cortex and was characterized by travelling waves in some lobules. Despite occurring only occasionally, the characteristic waves propagated from one end of a lobule to the other (i.e., along the mediolateral axis), which may contribute to the mediolateral-distance dependency of the CS synchrony (Figure S4A). We further examined how global dynamics changed as the anesthetic wore off. Under awake condition, overall synchrony was substantially decreased, while the phase reversal across the border of the olivocerebellar cross was still preserved well (Figure S4C). In addition, voluntary movements appeared to further perturb the synchrony.

### CS responses evoked by limb muscle stimulation

Using the large-scale optical measurements, we next examined the representation pattern of afferent (climbing fiber) inputs to the cerebellar cortex. For this purpose, we visualized CS responses elicited by the application of a weak and brief electrical pulse on a flexor muscle, and noticed that stimulation of a limb muscle caused some characteristic distribution of CS firing over the cortex (Figure 3). On the application of an electrical pulse on the left biceps brachii muscle (Figure 3A) of mouse #1032, the contrasting CS responses were observed in two segments (V5 and RP5) as exemplified in Figure 3B. In V5, the stimulation elicited a large positive transient, suggesting an increase in the number of CS-generating PCs within the segment. The transient signals (∼2 sec) were clearly detectable over resting signals that oscillated at 1.03 ± 0.22 Hz (n = 25) in individual trials. Averaging effectively emphasized the positive transients by offsetting the incoherent resting signals. In RP5, by contrast, resting oscillatory signals with a frequency of 1.07 ± 0.22 Hz (n = 25) were extinguished very transiently (∼1 sec) after stimulation, suggesting a decrease in the number of CS-generating PCs within the segment. A heatmap representing peak values of averaged FRET signals of individual segments, in which positive and negative responders (segments) are colored in red and blue, respectively (Figure 3C, top left), shows that a single muscle stimulus elicited CS responses that were widely distributed over the cortex almost symmetrically.

**Figure 3.**
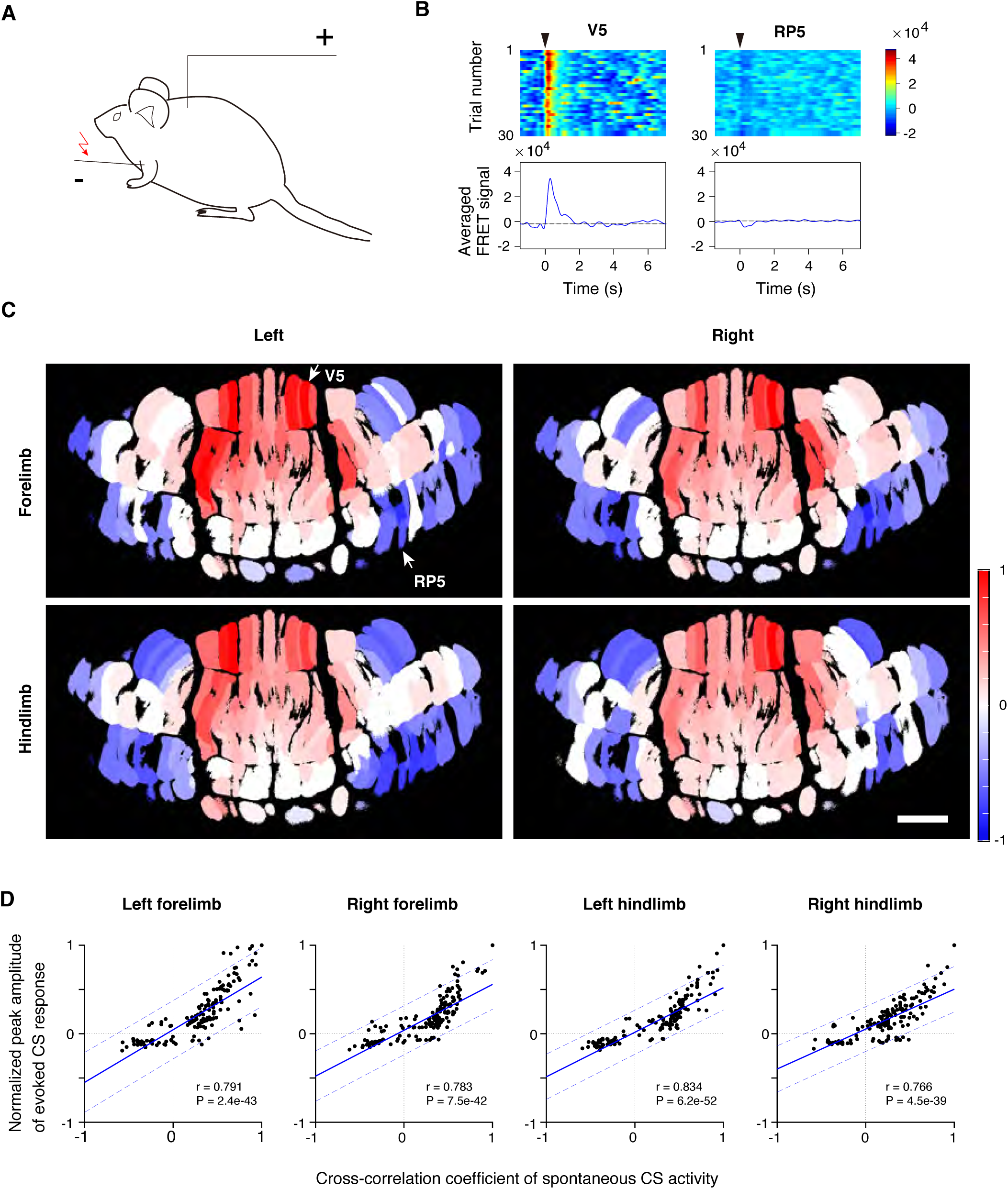
Similar distribution patterns of afferent inputs from four limb muscles over the cerebellar cortex. (A) Schematic drawing of a limb muscle stimulation experiment. Current pulses were delivered for 0.2 msec through an electrode inserted into the muscle. (B) Typical CS responses observed in two segments (V5 and RP5) with the electrical stimulation of biceps brachii of the left forelimb with a current pulse of 1.43 mA. Temporal profiles of FRET signals during 30 trials (top) and their averaged traces (bottom). (C) Heatmaps representing peak amplitude of the averaged CS responses (n = 30) with the stimulation of the biceps brachii of the left forelimb (1.43 mA) (top left) and right forelimb (1.47 mA) (top right), and biceps femoris of the left hindlimb (1.26 mA) (bottom left) and the right hindlimb (1.55 mA) (bottom right). Positive/negative responses (*i*.*e*. increase/decrease in the number of CS-generating PCs within the segment) were normalized against the maximum/minimum values and are colored in red/blue, respectively. Scale bar: 1 mm. (D) Relationship between spontaneous and evoked CS signals. For each of the four limb muscles, the segment that produced the maximum value of the average CS peak amplitude was identified, and evoked signals of individual segments are represented by their CS peak amplitudes normalized against the maximum value. Spontaneous signals of individual segments are represented by their cross-correlation coefficients of resting CS signals with the maximum value-producing segment as a reference. Pearson’s correlation coefficient (r) for linear regression and P value calculated for 196 data points are shown in each graph. Confidence intervals are shown by two blue broken lines. (A–D) All data are from L7-YC2.60 mouse #1032 (Table S1). See also Figure S5.

We then performed similar mapping with the electrical stimulation on each of the remaining three flexor muscles. Remarkably, although the stimulation was fairly specific to the respective targeted limb (Figure S5A), the four different inputs generated almost the same patterns (Figure 3C) independently of stimulation intensity (Figure S5B). These highly overlapping representation patterns were reproducibly observed in three other mice examined (Figure S5C). The overall CS activity patterns on the cortex suggest that excitatory CS responses were observed mainly within a cruciform area resembling the olivocerebellar cross while inhibitory responses were observed mainly in the four corners resembling the outside of the cross. The peak amplitude of evoked CS responses was compared with the cross-correlation coefficient of spontaneous CS activity for each segment (Figure 3D). Significant positive correlation (p < 0.01) indicates that the phase reversal generation of spontaneous CS activity (Figure 2E) shares a patterning mechanism with the positive/negative control of the firing probability of olivary neurons upon stimulation (Figure 3C). Taken altogether, these findings suggest that the olivocerebellar cross, originally delineated from spontaneous CS activity, is a functional division of the cerebellar cortex related to the responsiveness of olivary neurons to external stimuli.

The fraction of segments exhibiting excitatory and inhibitory responses was almost same for each muscle, irrespective of the stimulation intensity or the animal examined (Figure S5D). Interestingly, the latency of excitatory responses observed inside the cross was shorter than that of inhibitory responses outside the cross (Figure S5E), suggesting that direct spinal inputs mainly mediate the excitatory responses, while some indirect pathways contribute to the inhibitory responses. Minor fractions of CS responses, i.e., inhibitory responses inside the cross and excitatory responses outside the cross, exhibited a wide range of response latencies, suggesting the involvement of multiple neuronal pathways. In addition, we confirmed that every olivocerebellar segment, regardless of whether it was inside or outside the olivocerebellar cross, received inputs largely from all four limbs (Figure S5F).

### Bayesian decoding of CS population codes

The substantial overlap of segments responding to the four different peripheral stimuli (Figures 3C and S5C) suggested that the stimulation on a muscle would be represented as distributed CS population activities rather than the simple one-to-one correspondence between muscles and the cerebellum. To verify this possibility, we tried to decipher the CS population code by adopting probabilistic approaches. We carried out inference by applying Bayesian updating algorithm that was employed to infer animal position from ensemble firing patterns of hippocampal place cells (Zhang et al., 1998).

First, we estimated the stimulation onset time for the four limb muscles separately. Conditional probability density functions of evoked CS responses in each segment were directly estimated from observed FRET signals which exhibited considerable trial-by-trial variability (Figure S6). To combine the information for multiple segments, the likelihood was calculated from conditional probabilities of segmental FRET signals observed in each trial, and posterior probabilities were obtained given that a stimulation was applied (stimulation probability) or not applied (no stimulation probability) (Figure S7, see Methods for details). To examine the functional significance of the olivocerebellar cross, the likelihood was calculated for all segments (ALL) or segments inside and outside the cross (IN and OUT, respectively). For the inference of the stimulation onset time, we examined how these stimulation probabilities developed around the actual stimulation onset time (Figure 4). The stimulation probability for IN segments reached 1.0 in a manner similar to that for ALL segments, whereas the stimulation probability for OUT segments developed with a relatively large latency. It is thus assumed that IN and OUT segments contain different temporal information. In fact, although all of the stimuli were detectable by either IN or OUT segments when their intensity was sufficiently high (Figures S8A–S8C), OUT segments appeared to contribute to the high accuracy of the inference (Figure S8D–S8F). Considering the low frequency range of CS occurrence (∼1 Hz), we note that the difference between the actual and inferred onset times was only a few tens of milliseconds, strongly suggesting that the ensemble of CSs carries precise temporal information.

**Figure 4.**
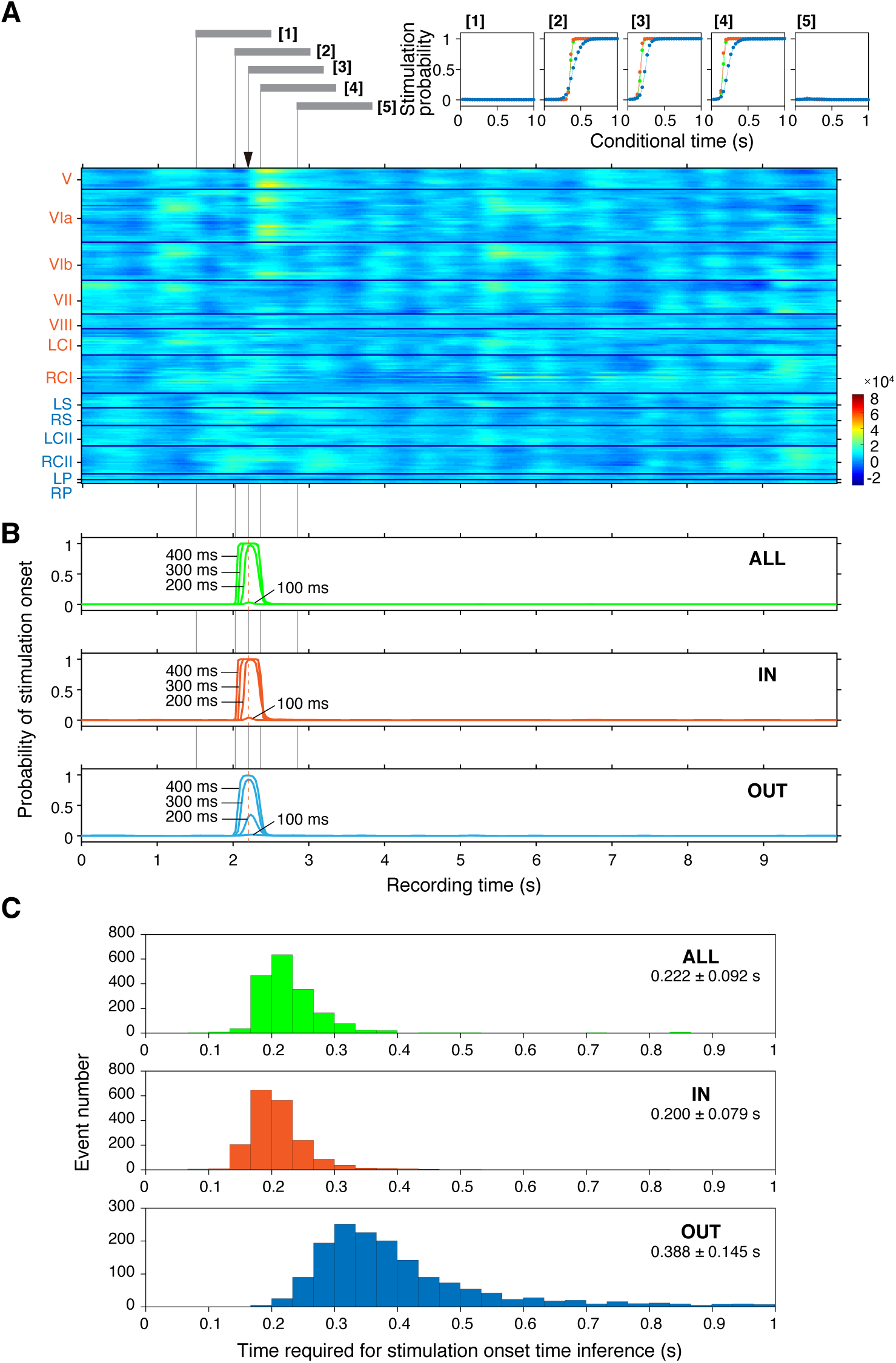
Bayesian inference of stimulation onset time from population activity, by single trial. (A) Time courses of color-coded FRET signals during a single-trial observation in all segments. At the recording time, indicated by an arrow, an electrical stimulus (duration: 0.2 msec, intensity: 1.83 mA) was applied to the biceps brachii of left forelimb. Segments are aligned as in Figure 2A. Lobule names are abbreviated and labeled as in Figure 2B. insets, The probabilities of the occurrence of an electrical stimulation at conditional time 0 (vertical lines) were computed using the likelihood values calculated from all segments (ALL, green), segments inside the olivocerebellar cross (IN, orange), or segments outside the cross (OUT, cyan). The time ranges of FRET signals used to calculate the stimulation probabilities are indicated by horizontal thick grey bars ([1] ∼ [5]). (B) The posterior probability (probability of stimulation onset) for the occurrence of the electrical stimulation (stimulation probability) was calculated for all recording time (from 0 to 10 s). Stimulation probabilities at 100, 200, 300 and 400 msec of conditional times are plotted against the recording time. Probability of stimulation onset was computed using likelihood values calculated from ALL (green), IN (orange), or OUT (cyan) segments. (A and B) Data were obtained from L7-YC2.60 mouse #1024. (C) Histograms of the conditional time at which the stimulation probability exceeded the control (no stimulation) probability for ALL (green), IN (orange), or OUT (cyan) segments. Data (mean ± s. d.) were calculated from 2,340 trials from 11 experiments from four L7-YC2.60 mice (Table S1). Each experiment had an equal number of trials for four limb muscles. See also Figures S6, S7, and S8.

Next, we designed experiments in which the four limb muscles were randomly stimulated, and attempted to infer the target muscle from CS ensembles. We performed the same statistics and Bayesian updating algorithm as above to obtain the time-dependent posterior probabilities of stimulation of four limb muscles each and no stimulation (Figure 5A). We found that the posterior probability for the target muscle progressively increased within 1 sec after the stimulation onset. We made an inference of the target muscle and no stimulation by selecting the maximal posterior probability for every time points (Table 1). On average, the rate of correct inference was 60% with the FRET data from ALL or IN segments but approximately 50% for OUT segments (Figure 5B and 5C). This decrease was not due to the small number of OUT segments (∼60 segments); even if the same number of segments were randomly chosen from ALL segments, they produced similar rates to ALL (Figure 5C). It must be added, however, that the correct inference rate was reduced when the segment number was lower than 20 (Figure 5D).

**Table 1.** Stimulation conditions and rates of correct inference.

| Mouse | Segment number |  | Stimulation intensity (mA) |  |  |  | Rate of correct inference* |  |  |
| --- | --- | --- | --- | --- | --- | --- | --- | --- | --- |
|  | IN | OUT | Left | Right | Left | Right | ALL | IN | OUT |
|  |  |  | forelimb | forelimb | hindlimb | hindlimb |  |  |  |
| #1024 | 148 | 55 | 0.64 | 0.41 | 0.63 | 0.69 | 0.583 | 0.694 | 0.444 |
|  |  |  | 0.94 | 0.70 | 0.88 | 0.98 | 0.528 | 0.417 | 0.389 |
|  |  |  | 1.23 | 0.98 | 1.11 | 1.26 | 0.556 | 0.583 | 0.528 |
|  |  |  | 1.53 | 1.27 | 1.35 | 1.55 | 0.861 | 0.889 | 0.500 |
|  |  |  | 1.83 | 1.56 | 1.85 | 1.85 | 0.694 | 0.722 | 0.417 |
| #1026 | 101 | 63 | 0.64 | 0.61 | 0.88 | 0.98 | 0.639 | 0.667 | 0.389 |
|  |  |  | 0.94 | 0.89 | 1.11 | 1.26 | 0.639 | 0.611 | 0.500 |
|  |  |  | 1.23 | 1.17 | 1.35 | 1.55 | 0.611 | 0.583 | 0.556 |
|  |  |  | 1.53 | 1.47 | 1.58 | 1.85 | 0.667 | 0.694 | 0.417 |
| #1034 | 169 | 69 | 0.64 | 0.70 | 0.80 | 0.31 | 0.611 | 0.667 | 0.417 |
|  |  |  | 0.94 | 0.98 | 1.03 | 0.59 | 0.472 | 0.500 | 0.278 |
|  |  |  |  |  |  | Mean | 0.624 | 0.639 | 0.439 |
|  |  |  |  |  |  | SD | 0.101 | 0.124 | 0.078 |

\*: Correct inference rates at 1 sec after the stimulation onset as shown in Figure 5C.

**Figure 5.**
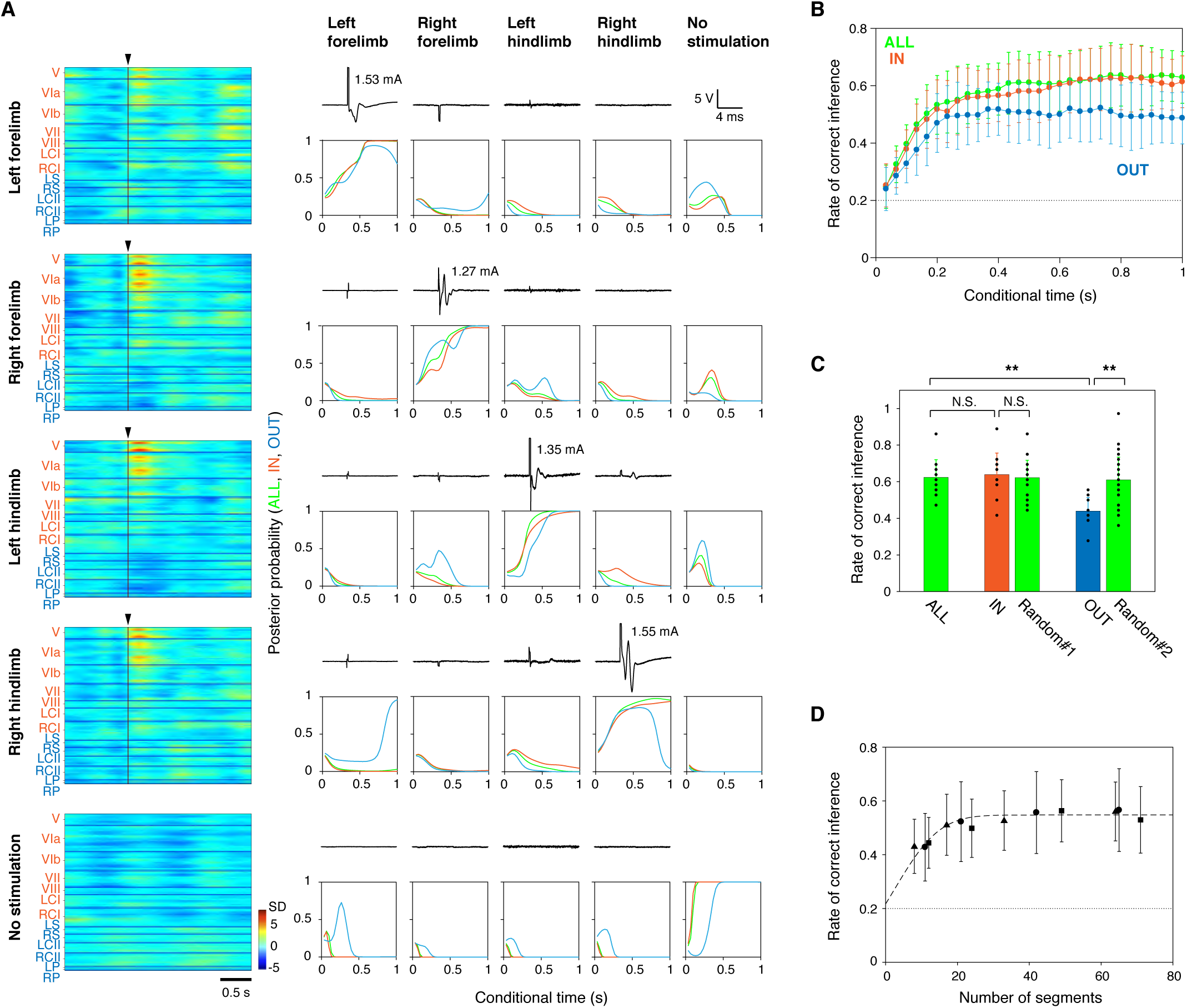
Bayesian inference of a stimulated muscle from ensemble CS firing patterns over the cerebellar cortex by single trial. (A) Representative population FRET measurements with muscle stimulation of left forelimb, right forelimb, left hindlimb, right hindlimb, or no limb (from top to bottom) and Bayesian inferences of their stimulation types. *left*, Time courses of color-coded normalized FRET signals in all segments. At conditional time = 0 (arrowhead), an electrical stimulus (duration: 0.2 msec) was applied. Segments are aligned as in Figure 2A. Lobule names are abbreviated and labeled as in Figure 2B. *right*, Electromyography traces (black lines) from −4 to +8 msec relative to the stimulation onset are shown for all four limb muscles. The posterior probabilities were calculated using the Bayesian updating algorithm with likelihoods of FRET signals observed in all olivocerebellar segments (ALL, green), segments inside the olivocerebellar cross (IN, orange), and outside the cross (OUT, cyan). This measurement was repeated 36 times with random choice of the five stimulation modes in each experiment. (B) The rate of correct inference of the stimulated muscle, including the control (no stimulation), is plotted against the conditional time for ALL, IN, or OUT. Data were obtained from 11 experiments using three L7-YC2.60 mice, and are shown as mean ± s. d. (n = 11) (Table 1). (C) Statistical data showing the correct inference rate at 1 sec after the stimulation onset for ALL (n = 11), IN (n = 11), and OUT (n =11). Random: the same number of segments were randomly selected from ALL as IN (#1) or OUT (#2). As three different selections were made, the sample size was threefold higher (n = 33). Data are shown as mean ± s. d. **: p < 0.01, Tukey-Kramer multiple comparison method, N.S.: not significant. (D) Relationship between the correct inference rate at 1 sec after the stimulation onset and the number of analyzed olivocerebellar segments. Segments were randomly selected from ALL in triplicate for each of the 11 experiments. Data are shown as mean ± s. d. (n = 33). The broken line indicates the normal cumulative distribution function (Figure S8). Data obtained from three different animals are shown with different symbols. (B and D) The dotted line indicates the chance level (0.2).

All these results suggest that the CS activities both inside and outside the olivocerebellar cross carry information about the timing and location of stimulation on a moment-to-moment basis.

## DISCUSSION

It has long been believed that microzones (Figure S1H) are the functional units of the cerebellum (Oscarsson, 1980; Voogd et al., 2013), as was restated by Ito (Ito, 1984) that “*… it is appropriate to define a corticonuclear microzone and an associated small group of nuclear cells which are dedicated to a single function of autonomic or somatic motor control*.”. Microzones have been assumed to work independently or coordinately, and this hypothesis could be tested by simultaneous recording from a large number of microzones. Due to technical limitations, however, no studies have been able to verify the hypothesis. For example, classical studies measured electrical activity from at most several tens of PCs in one experiment. Also, modern optical imaging techniques, such as two-photon excitation microscopy, covered no more than a few hundred PCs (equivalent to a few microzones) in a narrow field of view. Thus, brain-wide comprehensive visualization of PC activity over the cerebellar cortex is extremely challenging. To date, very few attempts have been made to examine CS activity across multiple lobules simultaneously. In this study, we assembled many technical trials to grasp the overall picture of the activation pattern of the olivocerebellar system. Among these methods was the combined use of the L7-Cre and tLSL-YC2.60 mouse lines, in which all the examined PCs were found to express YC2.60 abundantly. This comprehensive approach with the use of our custom-made macroscope allowed us to discover that the entire dorsal surface of the cerebellar cortex can be divided into approximately 200 segments, each of which has ∼100 PCs that fire CSs synchronously but at various incidence rates. Our large-scale optical measurements showed that each segment receives inputs mostly from multiple limbs (Figure S5F). More importantly, the measurements clearly showed that multiple olivocerebellar segments work coordinately, far from independently, to represent sensory signals originating from each of the four limbs. These findings should bring about a drastic change in our understanding about the operational principle of the cerebellum from a model based on parallel processing by a large number of independent local modules (Figure 6A) (Ito, 1984; Oscarsson, 1980) to one in which coordinated processing occurs with the olivocerebellar segments distributed over the entire cerebellar cortex (Figure 6B). Such a new understanding will force a global analysis of associative learning in the cerebellum. Climbing fiber inputs control the plasticity at parallel fiber-PC synapses; the occurrence of a CS in conjunction with parallel fiber inputs causes long-term depression (Ito, 1984), whereas the absence of CS occurrence allows the establishment of long-term potentiation (Dean et al., 2010). Although associative learning between mossy fiber inputs and climbing fiber inputs has been analyzed in individual microzones, the cortex-wide extension of an observation area will provide a more dynamic view of the cerebellar learning. Considering that the evoked CS activities were excitatory inside the olivocerebellar cross while inhibitory outside, for example, it is possible that the actual balance between potentiation and depression differs across the hierarchical cruciform structure.

**Figure 6.**
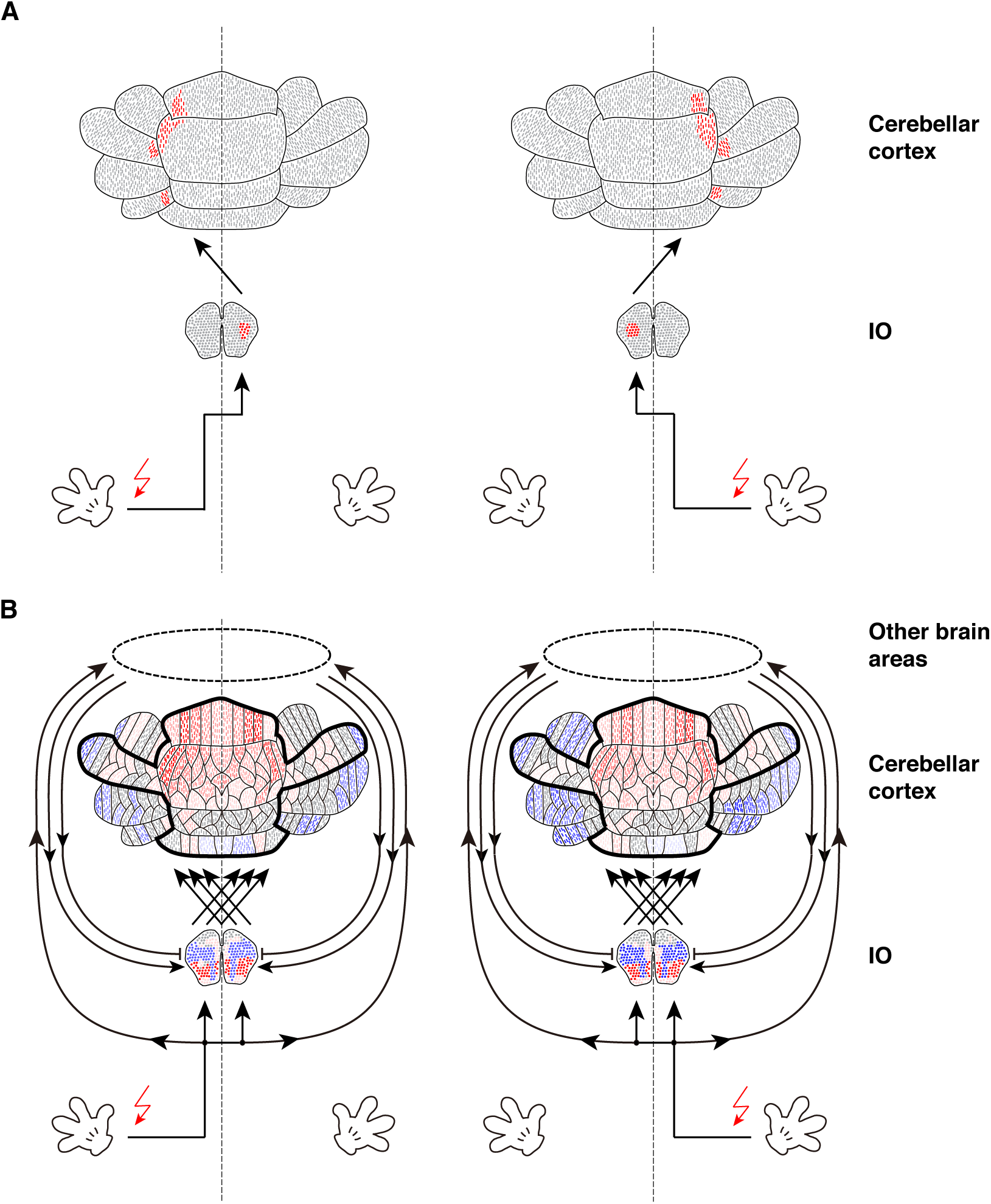
Cerebellar somatotopy hypothesis vs. distributed population coding. (A) Cerebellar somatotopy hypothesis. Neurons (principal neurons in the IO and PCs in the cerebellar cortex) exhibiting excitatory responses to limb muscle stimulation are shown in red. Non-responding neurons are shown in grey. Vertical broken lines indicate the midline. Sensory information is represented in CSs evoked in a population of PCs that are specific to the stimulated muscle. (B) Distributed population coding. The cerebellar cortex is divided into small compartments, i.e., olivocerebellar segments, and individual segments respond to external stimuli differently with trial-by-trial variability. Excitatory and inhibitory responses are indicated in red and blue, which reflect the net increase and decrease in the number of CSs, respectively, compared to the baseline activity in each segment, respectively. Color intensity reflects the firing probability of evoked CSs. In this model, sensory information is represented in ensemble patterns of CS activity of all segments, including non-responders (indicated in grey).

### Somatotopy in the cerebellar cortex

Sensations arising from the skin – such as the sense of touch, pressure, cold, heat, and pain – and from the muscles, tendons, and joints – such as the sense of the position and movement of limbs and trunk, effort, force and heaviness – are known as somatic sensations (Kandel et al., 2012). These multiple types of information are transmitted to the cerebrum and cerebellum separately via different ascending paths in the spinal cord. In the cerebral cortex, sensations from different body parts are first processed in different localized neuronal clusters in the postcentral gyrus, and this area is called the primary somatosensory cortex (S1) where somatotopic representation of the body parts has been thoroughly studied and well documented (Penfield and Rasmussen, 1950; Woolsey and Fairman, 1946). Earlier electrophysiological studies on the cerebellum assumed the presence of a somatotopic representation mechanism (Ekerot and Larson, 1979; Snider and Stowell, 1944). In this study, we uncovered a totally unique organization of the olivocerebellar system in which different muscle stimuli are encoded in similar ensemble patterns of CS activity of common segments over the cerebellar cortex (Figure 3C), and each segment is involved in processing information derived from multiple limb muscles (Figure S5F). These results strongly suggest that the IO transmits sensory information to the cerebellar cortex at every moment via different sets of climbing fibers and that the olivocerebellar system employs a distributed population coding scheme. The rate of electrophysiologically detected CSs in a single PC is so low (Andersson and Oscarsson, 1978; Catz et al., 2005; Keating and Thach, 1995; Kitazawa et al., 1998) that decoding CS time series data is considerably difficult. However, our new coding scheme ensures that multi-level gradations in each olivocerebellar segment and an assembly of multiple segments should contain a wealth of information.

We propose that multiple, possibly all, olivocerebellar segments coordinately work in space and time to represent peripheral states moment-by-moment, combining both excitatory and inhibitory CS responses with their characteristic time courses. The relative slowness of the inhibitory responses in OUT segments (Figures 4C and S5E) suggests the presence of relay neuronal circuits that innervate the IO (Figure 6B). The distributed population coding presented here is totally unique in that it operates in units of segments rather than neurons and that it implicates a cross-shaped architectural structure over the cortex. The newly discovered hierarchical structures are critical for describing the robust unified frame in the cerebellum to process a substantial amount of signals that are disseminated over the cortex in a democratic fashion.

### Probabilistic sensory representation in the cerebellum

By Bayesian analysis, we demonstrated that the segment-based distributed population coding scheme is truly effective. Despite considerable variability in evoked CS activity patterns among trials, we were able to infer correctly which limb muscle was stimulated from single-trial data. Our study demonstrated that a population of CSs observed in each olivocerebellar segment represented the conditional probability of the occurrence of various sensory events. A prominent feature of the cerebellar cortical circuits is their clear input-output relationship (Figure S1A); since the publication of the cerebellum theory by David Marr more than 50 years ago (Marr, 1969), quite a few ideas have been proposed regarding the cerebellar computation (Table S3). However, we are the first to regard a probabilistic representation as the function of climbing fiber inputs.

Helmholtz originally proposed that perceptions are extracted from sensory input data by probabilistic modelling of their causes (Von Helmholtz, 2005). The predictive coding theory (Rao and Ballard, 1999) has stated that sensory signals are compared with internally generated top-down predictions, and differences between these two quantities, i.e., prediction errors, are further processed in a hierarchy of networks. Friston further extended the theory using statistics signals based on the Bayesian brain hypothesis (Knill and Pouget, 2004) that perceptions are the results of Bayesian inversion of a causal model, which is updated by processing of sensory signals according to Bayes theorem. The free-energy principle (Friston, 2005) formulates how neuronal processing could minimize variational free energy, either by changing sensory inputs via action or by updating internal models (predictions) via perception. Although the free-energy principle is expected to provide a unified theory of the brain, integrating data and theory relating to action, perception, and learning (Friston, 2010), its biological implementation in real neural networks remains elusive. The probabilistic representation of sensory events by a population of CSs elucidated in our study, remarkably, can be well described in the framework of the free-energy principle. Because sensory processing is inseparable from the generation of motor commands in the framework, further analysis of information processing operated in units of the olivocerebellar segments will provide a better understanding of how the cerebellum uses statistics of sensory signals to control movements and perception.

## SUPPLEMENTAL INFORMATION

Supplemental Information includes eight figures and three tables.

## ACKNOWLEDGMENTS

The authors thank Drs. Michael Häusser, Beverley Clark, Michael London, Jesper Sjöström, Taro Ishikawa, Kazuo Kitamura, Benjamin Judkewitz, Matteo Rizzi and Jenny Davie for help with performing electrophysiology and in vivo two-photon imaging; Dr. Susumu Tonegawa and Chie Sano for providing L7-Cre transgenic mice; Yumi Sato for help with preparing adenovirus; Dr. Hiroshi Kurokawa for help with programming; Dr. Naomi Kogo for help with immunohistochemistry; Dr. Izumi Sugihara for valuable comments on the olivary neuronal projection; Drs. Masami Tatsuno, Kazuhiko Yamaguchi and Takeru Honda for fruitful discussion; Drs. Shigeyoshi Fujisawa, Taro Toyoizumi and Lukasz Kusmierz for critical reading of the manuscript. This work was supported in part by grants from the Japan Ministry of Education, Culture, Sports, Science, and Technology Grant-in-Aid for Scientific Research on Innovative Areas: Resonance Bio (15H05948), the Brain Mapping by Integrated Neurotechnologies for Disease Studies (Brain/MINDS), and Integrated Symbiology (iSYM) research program.

## AUTHOR CONTRIBUTIONS

T.M. and A.M. conceived the whole study. T.M. designed and performed all the experiments, analyzed the data for preparation of the manuscript. T.Y. provided knowledge and expertise in electronics and software for constructing and operating the macroscopy system. S.I., S.Kuroki and A.M. prepared YC2.60-transgenic mouse lines for studying neural circuits. T.I. and S.Kakei contributed to muscle stimulation experiments. A.M. and T.M. wrote the manuscript. A.M. supervised the whole project. All authors contributed to the final manuscript.

## DECLARATION OF INTERESTS

The authors declare no competing interests.

## Methods

### Animal preparation

All experimental procedures were performed in accordance with the guideline of the Animal Experiment Committee of RIKEN. A mixture of fentanyl (0.05 mg/kg), midazolam (5.0 mg/kg), and medetomidine (0.5 mg/kg) was used to anesthetize mice according to the method described previously (Mrsic-Flogel et al., 2007). All the animals used in this study are listed in Table S1.

### Expression of Ca^2+^ indicator proteins in cerebellar PCs

Recombinant adenovirus carrying YC2.60 was injected into the lateral ventricle of an ICR mouse embryo at embryonic day 11 as described previously (Yamada et al., 2011). Mice expressing YC2.60 were identified during postnatal days 1–3 by viewing their transcranial fluorescence. L7-Cre transgenic mice (provided from Dr. Susumu Tonegawa) were cross-bred with tLSL-YC2.60 mice (Kuroki et al., 2018) to obtain L7-YC2.60 mice.

### Immunohistochemistry

Immunohistochemistry was performed according to the method described elsewhere (Yasuda et al., 2017). Mouse monoclonal antibody against calbindin D28k (#300) was purchased from Swant and rabbit polyclonal antibody against GFP (MBL598) was purchased from MBL. Alexa Fluor 594-conjugated anti-mouse IgG (A11034) and Alexa Fluor 488-conjugated anti-rabbit IgG (A11032) were purchased from Molecular Probes. Images were acquired with an inverted laser scanning confocal microscopy system (FV1000, Olympus) equipped with 473 nm and 559 nm lasers, an excitation dichroic mirror (DM405/473/559), an emission dichroic mirror (SDM560), two emission filters (BA490-540 and BA575-675) and an oil-immersion objective (UPL SAPO 20× NA:0.75, Olympus).

### Surgery

A custom-made stainless-steel head plate was attached to the mouse skull with surgical cyanoacrylate and dental cement (Super-bond C&B, Sun Medical) under anesthesia. After at least one week, a section of the skull was removed along with the dura mater under anesthesia. For electrophysiological measurements, a part of the cerebellar cortex was covered with 2% agarose and a 1-mm thick cover glass (Matsunami). For chronic observation experiments, a tightly sealed cranial window was constructed using a 1-mm thick cover glass (8 mm × 5 mm, or 8 mm × 4 mm) (Matsunami).

### *In vivo* Ca^2+^ imaging combined with electrophysiology

Anesthetized head-fixed mice on postnatal day 20–220 were used. Ca^2+^ (YC2.60) signals in the dendrites of PCs were acquired in a frame-scan mode (256 × 256 pixel, 2–20 Hz) using an upright two-photon laser-scanning microscope (LSM710MP, Carl Zeiss) equipped with a 20× water-immersion objective (W Plan-Apochromat 20×/1.0 DIC D=0.17 M27 70mm, Carl Zeiss). Image acquisition at greater than 8 Hz was carried out using the line-step mode of the ZEN2010 software (Carl Zeiss) with a pair of Galvano scanners. The Ti:sapphire laser (Chameleon Ultra, Coherent) for two-photon excitation was tuned to 820 nm. Emitted fluorescence was short-pass filtered (630 nm), split with a dichroic mirror (510 nm), band-pass filtered (460–500 nm and 525–560 nm for cyan and yellow fluorescence, respectively) and detected using GaAsP (gallium arsenide phosphide) detectors. Extracellular single-unit recording was carried out on the optically imaged PCs in lobule V using borosilicate glass pipettes (1–3 MΩ) filled with an extracellular solution containing 150 mM NaCl, 2.5 mM KCl, 10 mM HEPES, 2 mM CaCl_2_, 1 mM MgCl_2_ (pH 7.3, ∼300 mOsm/L). Body temperature was monitored with a rectal thermistor and maintained at 36–37°C using a heating pad. Electrophysiological signals were acquired at 100 kHz using MultiClamp 700B (Molecular Devices) and Digidata 1440A (Molecular Devices) controlled by pClamp10 (Molecular Devices). The electrophysiological recording and two-photon imaging were synchronized by a trigger pulse generated by the image acquisition software ZEN2010. CSs were identified from extracellular single-unit records by using a hierarchical clustering method (T.M., *et al*., in preparation).

### Ultra-wide field Ca^2+^ imaging

*In vivo* imaging of the entire dorsal surface of the mouse cerebellar cortex under anesthetized or awake conditions was carried out using a custom-made tandem-lens macroscope as described elsewhere (Kuroki et al., 2018).

### Stimulation and electromyography recording of biceps muscles

Electrical stimuli to limb muscles (biceps brachii of the forelimb of biceps femoris of the hindlimb) were carried out using stimulus isolators (ISO-Flex, A.M.P.I.) regulated by an Arduino microcontroller. The device monitored trigger pulses delivered from the CCD camera at the onset of acquisition of every image to adjust the stimulation timing. A brief electrical stimulus (0.2 msec) was delivered synchronously at the onset of image acquisition (exposure time: 33.3 msec) (see below). A thin stainless-steel wire (SUS304, Unique Medical) inserted into a muscle was used as an electrode for stimulation. Electromyography of a limb muscle was recorded at 100 kHz using the pClamp10 software (Molecular Devices) with a multichannel amplifier (AB-611J, Nihon Kohden) and a pair of recording electrodes (SUS304, Unique Medical) inserted into the biceps muscle.

### Data analysis

All of the following data analyses were performed using custom-made software written in MATALB.

### Preprocess

Image registration of ECFP and Venus fluorescence signals of YC2.60 was carried out using a combination of a rotation-invariant phase-only correlation technique for size/angle adjustment, and a phase-only correlation technique for x-y shift adjustment at the subpixel level (Takita et al., 2003). These registrations were performed for the two emission (ECFP and Venus) channels first individually to compensate for motion artifacts and then in pairs for calculation of ratios. After correction for photobleaching in the two channels, mean-subtracted ratio images, ℝ(*t*), were constructed as follows,

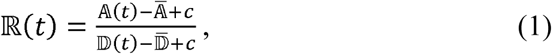

where 𝔻(*t*) is the photobleaching-corrected image of the donor (ECFP) channel at time *t*, 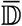 is the temporal mean of donor images, 𝔸(*t*) is the photobleaching-corrected image of the acceptor (Venus) channel at time *t*, 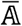 is the temporal mean of acceptor images, and *c* is the offset value used to prevent division by zero. After converting mean-subtracted ratio images from 32-bit floating-point numbers into a 16-bit resolution with a fixed range throughout the experiment, ratio images were constructed for the following analyses. In mean-subtracted ratio images, the background fluorescence signals outside cells settled down to a substantial level with negligible variance throughout the recording period. This is a marked contrast with conventional ratio images, in which ratio values of background pixels exhibit large variations due to very small values of both the numerator and denominator. Mean-subtracted ratio images were used for the independent component analysis (see below). “FRET signals” correspond to mean-subtracted ratio values with a 16-bit resolution. Mean-subtracted ratio signals normalized against their standard deviation are called as “normalized FRET signals”.

### Automatic identification of segments

After dimensional reduction using the principal component analysis, the independent component analysis (Mukamel et al., 2009) was used for the identification of PC clusters that exhibit synchronous CS firing from data that typically contained 6-min long continuous mean-subtracted ratio images. After the identification of independent components, continuous PC clusters greater than 15,820 μm^2^ (500 pixels) were segmented from spatial filters. All identified segments were averaged in shape and assigned on the cerebellar cortical map of each mouse examined.

We found that the central segment(s) in lobule V extended into lobule VIa (Figure S2I). Because this feature is observed reproducibly in all mice examined in this study, we used the center of these extended segments to draw the midline on the cerebellar cortex. Segments are numbered according to their mediolateral distance from the midline (Figures 1D, S2I and S3C). Within the vermis, segments are numbered positively to the right and negatively to the left.

### Phase analysis of FRET signals observed within segments

A bandpass filter (1.2 ± 0.3 Hz) was constructed using the MATLAB function *firls*. Its application to FRET signals for zero-phase digital filtering was performed using the MATLAB function *filtfilt*. A phase value was calculated from bandpass filtered FRET signals using a Hilbert transform. The mean phase value of a population of FRET signals from multiple segments was calculated as the angle of the mean vector, Θ, using the following equation,

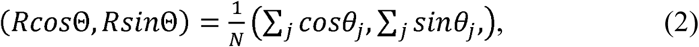

where *R* is the length of the mean vector (0 ≤ R ≤ 1), *N* is the number of segments and θ is the phase of segmental FRET signals. *R* is regarded as an index of the degree of the synchrony of FRET signals among segments since the variance of the phase of a population of FRET signals, *V*, is expressed as,

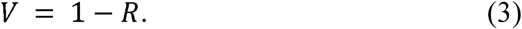

The frequency of spontaneous CS activity observed in each olivocerebellar segment was calculated from multiple 1-minute measurements (26, 19, 25 and 12 measurements for mice #1024, # 1026, #1032 and #1034, respectively).

### Bayesian inference of the stimulation onset time from a population activity of CSs

The minimal intensity of the injected current required for limb movement was determined for each of the four limb muscles (biceps brachii of left and right forelimbs and biceps femoris of left and right hindlimbs) under anesthesia. FRET signals of all segments were measured when one of the four limb muscles, *s*, was stimulated with a current intensity, *a*, greater than the minimal intensity. The stimulation was repeated 30 times with intervals of 10 ± 0.2 sec.

By using the data acquired from m trials, we constructed histograms of FRET signals, 𝕏_*i*_(*t*) {*x*_*i*,1_(*t*),…,*x*_*i,m*_(*t*)}, observed within a segment *i* at a conditional time *t*_*c*_ relative to the stimulation onset,

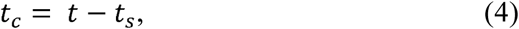

where *t* is the recording time when the FRET signal *x* was acquired and *t*_*s*_ is the stimulation onset time. Because FRET signals were acquired at 30 Hz, *t* is a discrete variable with a constant interval of 33.3 msec, Δt. The stimulation onset was exactly synchronized with the onset of FRET signal acquisition, and therefore *t* and *t*_*c*_ shared the same Δt (33.3 msec). For instance, FRET signals observed in segment 5 in lobule V (V5) (Figure S6A) and segment 12 in left crus II (LCII12) (Figure S6E) are shown in the top panel in Figures S6B and S6F, respectively. A single current pulse (0.2 msec, 1.83 mA) was applied to the biceps brachii of the left forelimb at time 0 and the stimulation was repeated 30 times with intervals of 10 ± 0.2 sec. Evoked FRET signals varied from trial to trial but exhibited a characteristic distribution over time (black horizontal bars in Figures S6C and S6G). Gaussian parameters of the mean and variance, 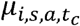 and 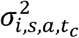, respectively, were estimated by fitting a Gaussian function,

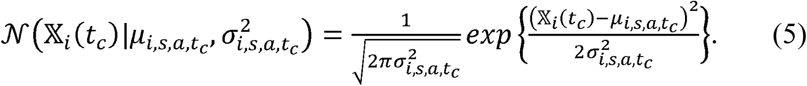

to the distribution of FRET signals, 𝕏_*i*_(*t*), at each conditional time, *t*_*c*_. The estimated probability density functions are arrayed in Figures S6C and S6G.

The conditional probability, *P*_*i*_(*x*_*i*_(*t*)| *s,a,t*_*c*_), of observing FRET signals, *x*_*i*_(*t*), in segment *i* at time *t* given the stimulation of muscle *s* with a current intensity of *a* at time -*t*_*c*_ were calculated by using the above-mentioned Gaussian parameters as follows,

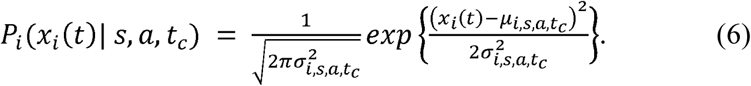

Despite the uniformity of the stimulus intensity, the conditional probability was variable, reflecting trial-by-trial variation in evoked FRET signals (Figures S6D and S6H).

To obtain the likelihood, we basically calculated the conditional probability during the conditional time *t*_*c*_ from +33 msec to 1 sec. The likelihood of observing 𝕏 = {*x*_1_,…,*x*_*i*_,… *x*_*n*_}of *n* segments at time *t* given the stimulation condition {*s,a,t*_*c*_} was calculated as follows,

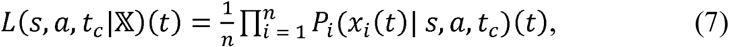

where *x*_*i*_(*t*) is the FRET signal observed in segment *i* at time *t* and *P*_*i*_(*x*_*i*_(*t*)| *s,a,t*_*c*_) is the conditional probability calculated according to Eq. 6.

For the calculation of the baseline probability, the FRET signals observed in segment *i* during a period of 2 sec before the onset of muscle stimulations in all trials were pooled, and a Gaussian function,

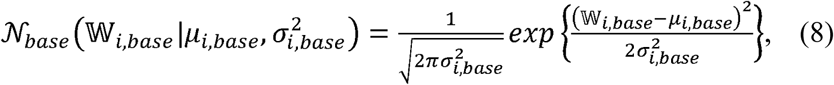

was fit to the distribution of the pooled signals, 𝕎_*i,base*_. Here, *μ*_*i,base*_ is the mean and 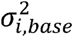 is the variance of the baseline FRET signals. The conditional probability, *P*_*i*_(*x*_*i*_(*t*)| *base*), of the single-trial FRET signals, *x*_*i*_(*t*), observed in segment *i* at time *t* given no muscle stimulation was calculated by using the Gaussian parameters estimated as described above, as follows:

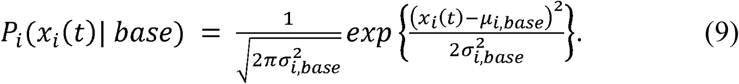

The likelihood of observing *x* = {*x*_1_,…,*x*_*i*_,…*x*_*n*_} of *n* olivocerebellar segments at time *t* given no muscle stimulation was calculated as follows:

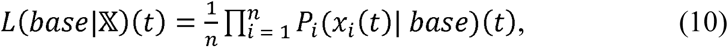

where *x*_*i*_(*t*) is FRET signals observed in segment *i* at time *t* and *P*_*i*_(*x*_*i*_(*t*)| *base)* is the conditional probability calculated according to Eq. 9.

Estimation of the stimulation onset time from single-trial population FRET signals was carried out using a Bayesian updating algorithm:

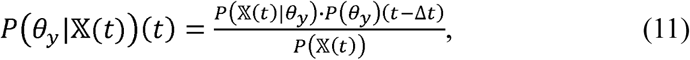

where 𝕏(*t*) = {*x*_*1*_(*t*),…, *x*_*n*_(*t*)} are the FRET signals observed in *n* segments at time *t, θ*_*y*_ indicate whether electrical stimulation to a limb muscle does not occur (y = 0) or occurr (*y* = 1) at conditional time 0, *P*(*θ*_*y*_)(*t*-Δ*t*)is the prior probability of the occurrence (*y* = 1) or nonoccurrence (*y* = 0) of the muscle stimulation at time *t*-Δ*t, P*(𝕏(*t*)) is the marginal probability of the observation of 𝕏(*t*), *P*(𝕏(*t*) |*θ*_*y*_) is the likelihood of the observation of 𝕏(*t*) given *θ*_*y*_ calculated according to Eq. 7 and Eq. 10), and *P*(*θ*_*y*_ |𝕏(*t*))(*t*)is a posterior probability of the stimulation condition *θ*_*y*_ at time *t* given 𝕏(*t*). Initially, we used the actual probabilities of the application of the electrical stimulation as prior probabilities; 299/300 for nonoccurrence and 1/299 for the occurrence of the muscle stimulation because an electrical pulse was delivered at only one time point within a single trial experiment, which had 300 time points (= 10 sec). Then, the posterior probabilities were calculated recursively using the previous posterior probability as the prior probability in the next step.

An example of the calculation of the posterior probability from single-trial experiment is shown (Figure S7). Conditional probabilities for stimulus occurrence and nonoccurrence were calculated for FRET signals observed in all segments. Probability density functions (normalized to the maximum value) given occurrence and nonoccurrence of muscle stimulation are shown in red and blue, respectively, for each segment; six segments are exemplified (Figure S7B). The functions for occurrence were conditional time-dependent, while the functions for nonoccurrence were time-invariant. FRET signals are shown by black dots, and their conditional probabilities are shown as vertical lines. The likelihoods of observing the FRET signals given muscle stimulation (red) and no muscle stimulation (blue) at various conditional times are shown in Figure S7C. These likelihood values were used to compute posterior probabilities according to the Bayesian updating algorithm. In this trial, the posterior probability for stimulation occurrence surpassed that for nonoccurrence between 233 and 267 msec (Figure S7D).

For the inference of the stimulation onset time, we computed the posterior probability for 1 sec starting from all the 300 time points within each trial. Examples of the posterior probability calculated from five different time periods of single-trial FRET signals are shown in Figure 4A. If *P*(*θ*_1_ |𝕏(*t*))(*t*) became greater than *P*(*θ*_0_ |𝕏(*t*))(*t*), *i*.*e. P*(*θ*_1_ |𝕏(*t*))(*t*) > 0.5, the stimulation onset time of the trial was estimated as follows. The posterior probability of stimulation occurrence calculated from the actual onset time, such as [3] in Figure 4A, most quickly reached 1. However, the posterior probability could reach 1 even when the calculation was started from time points around the actual onset time, as long as the recursive calculation continued for longer conditional times. Therefore, we identified peaks of posterior probability curves, which were plotted against the starting time point (Figure 4B), at earlier conditional times in which fewer than three time points with a probability of 1 were found. A mean value of the time points showing peak values of the posterior probability curves was used as an estimate of the stimulation onset time. If the difference between the estimated and actual onset times was less than 0.5 sec, we considered that the onset time was correctly inferred.

### Bayesian inference of a target muscle from a population activity of CSs

Estimation of the target muscle from single-trial population FRET signals was also carried out with the Bayesian updating algorithm. For this analysis, we performed the following measurements. First, one of the four limb muscles was stimulated 30 times with a fixed amount of current at random intervals of 10 ± 0.2 sec. The measurements of FRET signals evoked by muscle stimulation were performed for all four limb muscles separately with appropriate amounts of current adjusted for each muscle. These data were used for the estimation of Gaussian parameters of the time-dependent probability distribution function (Eq. 5) of FRET signals of individual segments. Then, current intensities were fixed for each muscle and one of the four limb muscles was randomly stimulated using four stimulation isolators controlled with an Arduino microprocessor. Random stimulation, including the no stimulation control, was repeated a total of 36 times with random intervals of 10 ± 0.2 sec. Electromyography was also recorded at 100 kHz using pClamp10 software. Trigger signals delivered from the Arduino were also recorded at 100 kHz and used to determine the target muscle and the time of stimulation of each trial. By using various current intensity sets, the measurements of FRET signals were repeated for three mice (Table 1). From these data with random stimulation, we calculated the likelihoods for all four muscles and no stimulation control using the predefined Gaussian parameters, and then applied them to the Bayesian updating algorithm (Eq. 11). In this case, *θ*_0_ is the no stimulation control, *θ*_1_ is the stimulation of biceps brachii of the left forelimb, *θ*_2_ is the stimulation of biceps brachii of the right forelimb, *θ*_3_ is the stimulation of biceps femoris of the left hindlimb, *θ*_4_ is the stimulation of biceps femoris of the right hindlimb. Actual probabilities of five stimulation modes (left forelimb: 0.23, right forelimb: 0.20, left hindlimb: 0.18, right hindlimb: 0.22, no stimulation control: 0.17) were used as the initial values of the prior probability. For the estimation of the target muscle, the stimulation onset time *t*_*s*_ was given. The maximum a posteriori probability (MAP) estimate was used to infer the stimulation mode as follows:

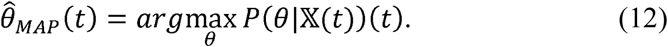

### Statistics

No statistical methods were used to predetermine sample size. Correlations were Pearson correlation coefficients with P values calculated after Fisher’s z-transformation. Ninety-five per cent confidence intervals with lower and upper confidence limits are given for each comparison. For multiple comparison, the Tukey-Kramer method and one-way ANOVA with post-hoc tests were applied. The equality of probability density functions was measured using Kolmogorov–Smirnov test. P□< □0.05 was considered statistically significant. For multiple comparisons, Bonferroni adjustment was used to adjust the significance level.

## SUPPLEMENTARY FIGURE LEGENDS

**Figure S1.**
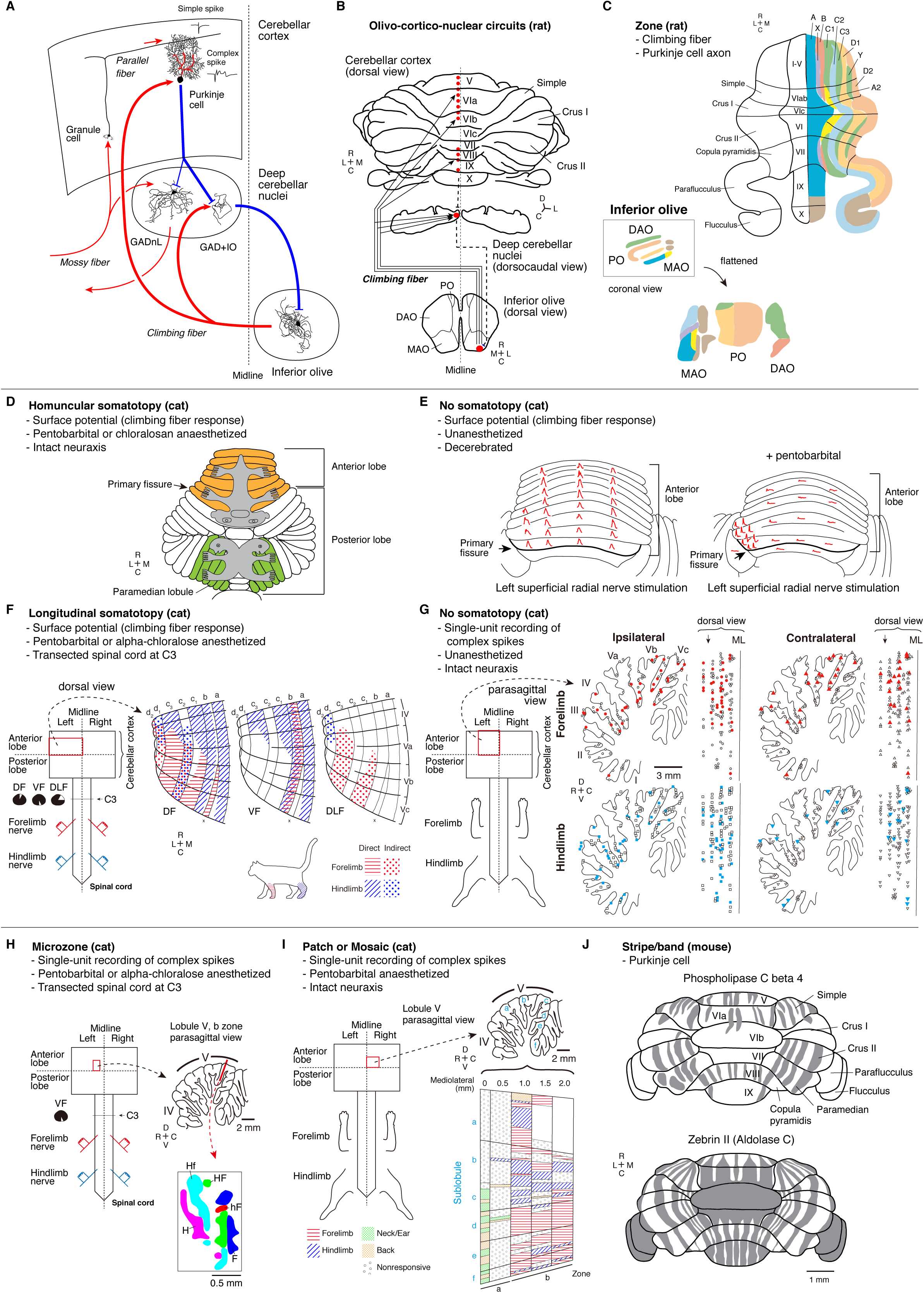
Cerebellar circuits and cortical compartments. (A) Schematic drawing of the main circuits in the cerebellum. Olivo-cortico-nuclear circuits are indicated by bold lines. Excitatory and inhibitory inputs are shown in red and blue, respectively. GADnL: glutamic acid decarboxylase (GAD)-negative non-GABAergic large projection neuron (Uusisaari and Knopfel, 2011). GAD+IO: GAD-positive GABAergic neuron projecting to the inferior olive (Uusisaari and Knopfel, 2011). Simple spikes are produced endogenously in PCs, and their firing pattern is modulated by excitatory synaptic inputs from parallel fibers that relay various inputs delivered from mossy fibers. A complex spike (CS) is evoked by a climbing fiber input originated from the contralateral inferior olive (IO). (B) Olivo-cortico-nuclear circuits in the rat brain (Sugihara and Shinoda, 2004). Climbing fibers from a region of the inferior olive contralaterally terminate in a narrow parasagittal array of PCs across transverse lobules (Sugihara and Shinoda, 2004). The climbing fibers also produce collateral branches that innervate a restricted region of a contralateral deep cerebellar nucleus, which is postulated to receive inhibitory inputs from the PCs (Sugihara et al., 2009) and send inhibitory outputs to the same IO region (Chaumont et al., 2013; Ruigrok and Voogd, 1990). Therefore, the olivocerebellar system is considered to comprise many closed-loop modules composed of olivo-cortico-nuclear connections. DAO: dorsal accessory olive. MAO: medial accessory olive. PO: principal olive. (C) Parasagittal zones (A, A2, B, C1, C2, C3, D1, D2, X and Y) were defined according to the anatomical arrangements of climbing fibers and PC axons (Voogd et al., 2013), as shown in Figure S1B. DAO: dorsal accessory olive. MAO: medial accessory olive. PO: principal olive. Modified from (Voogd et al., 2013). (D**–**G) The pros and cons of the cerebellar somatotopy hypotheses. (D) Homuncular somatotopic maps proposed in the cat cerebellar cortex (Snider, 1958; Snider and Stowell, 1944). Surface potentials were recorded under pentobarbital- or chloralosan-anesthesia. Projecting areas from the touch sense organs and from the proprioceptive endings that monitor muscle behavior are schematically depicted in grey on the flattened cerebellar cortex. Modified from (Snider, 1958). (E) The absence of somatotopy in the cerebellum in decerebrated unanesthetized cats (left). Localized responses were observed only under pentobarbital anesthesia (right). Modified from (Combs, 1954). (F) Somatotopic organization of the termination zones of spino-olivocerebellar paths. The parts of lobules IV and V in the left half of the cerebellar cortex are shown. The projection from the dorsal funiculus to c2 zone and the projection from the ventral funiculus to b zone are bilateral, whereas all other projections are ipsilateral. Surface potentials evoked by electrical stimuli of peripheral nerves were recorded in pentobarbital- or alpha chloralose-anesthetized cats with the spinal cord transected at the C3 level with examined funiculus maintained: dorsal (DF), ventral (VF) or dorsolateral (DLF) funiculus. Modified from (Ekerot and Larson, 1979). Anatomically defined zones are labeled with uppercase letters (C), whereas electrophysiologically defined zones are labeled with lowercase letters (F). (G) The absence of somatotopy in unanesthetized cats with intact neuraxis. Distribution of PCs in the cerebellar anterior lobe with (filled symbols) or without (open symbols) the climbing fiber inputs from cutaneous mechanoreceptors of the ipsilateral forelimb (circle), ipsilateral hindlimb (square), contralateral forelimb (triangle) and contralateral hindlimb (inverted triangle) (left). Mediolateral distributions of the cells in the recording area left from the midline (ML) are shown in the right. The arrow marks the border-line between vermis and pars intermedia at the surface of lobule Vb. Modified from (Leicht and Schmidt, 1977). (H**–**J) Compartmentalization of the cerebellar cortex. (H) The organization of microzones in the cat cerebellar cortex. Microzones were defined by single-unit recordings in b zone within lobule V on the basis of complex spike (CS) responses elicited with direct stimuli of ipsilateral or contralateral ulnar (forelimb) or sciatic (hindlimb) nerves (Andersson and Oscarsson, 1978). The track of a microelectrode used for recording is shown in a red line in a parasagittal section. H: activation exclusively from hindlimb nerves. Hf: short-latency activation from hindlimb nerves and long-latency activation from forelimb nerves. HF: short-latency activation from hindlimb and forelimb nerves. hF: short-latency activation from forelimb nerves and long-latency activation from hindlimb nerves. F: activation exclusively from forelimb nerves. Single PC activities were recorded extracellularly in pentobarbital- or alpha chloralose-anesthetized cats. The spinal cord was transected at the C3 level except for VF (ventral funiculus). Defined zones are labeled with lowercase letters. Modified from (Andersson and Oscarsson, 1978). (I) A diagram of an unfolded vermal cortex of lobule V of the cat. The major body areas whose cutaneous stimulation elicited climbing fiber responses emerged in a patchy pattern. Modified from (Robertson, et al., 1982). (J) The organization of stripes and bands in the mouse cerebellar cortex. These compartments were defined histochemically on the basis of gene expression in PCs (Apps and Hawkes, 2009). Areas with expression of phospholipase C beta4 (top) and aldolase C (bottom) are shaded. Modified from (Apps and Hawkes, 2009). (B, C, F**–**J) C: caudal. R: rostral. D: dorsal. V: ventral. L: lateral. M: medial.

**Figure S2.**
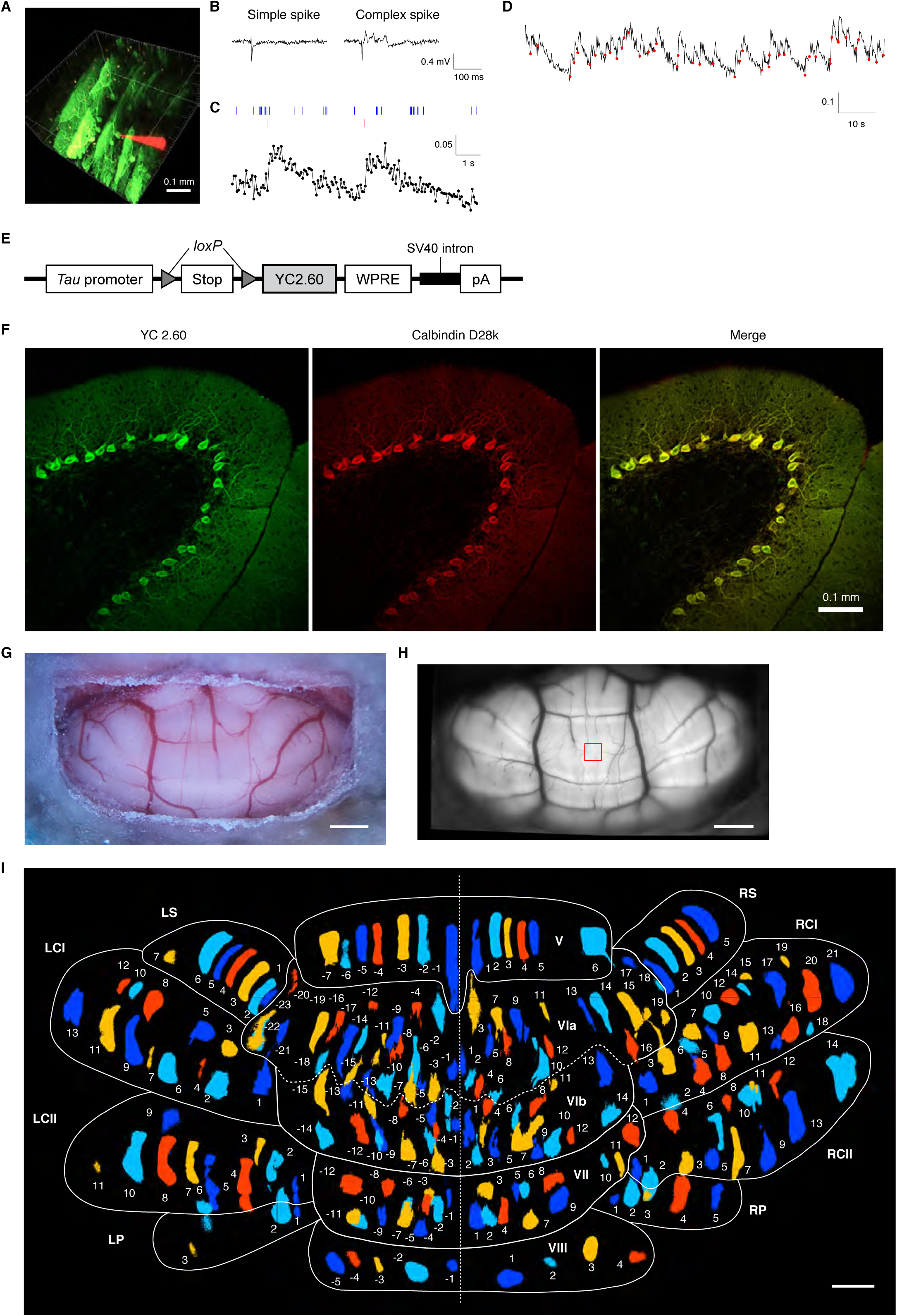
Detection of complex spikes in PCs from dendritic signals of YC2.60 for compartmentalization of the mouse cerebellar cortex. (A) Venus fluorescence (green) from YC2.60-expressing PCs and Alexa Fluor 568 fluorescence (red) in a glass pipette used for extracellular recording. (B) Typical waveforms of an extracellular single-unit recording from a PC that contained a simple spike and a complex spike (CS). Signals were acquired at 100 kHz. (C) A time course of normalized FRET signals sampled at 16 Hz with the occurrence of simple spikes (vertical blue lines) and CSs (vertical red lines). (D) A similar but extended time course of normalized FRET signals with actual CSs (filled red circles) that were electrophysiologically identified. It was concluded that the YC2.60 readout can tell the timing of CS occurrence faithfully. (A–D) Experiments were performed using an anesthetized ICR mouse, which had been infected with a recombinant adenovirus expressing YC2.60. The virus-mediated gene transfer method achieved sparse, random labeling of PCs, allowing us to examine calcium signals at the single cell level. (E) Structure of the tLSL-YC2.60 transgene (Kuroki et al., 2018). (F) Immunohistochemical localizations of ECFP/Venus (YC2.60) and a PC-marker (calbindin D28k) in the cerebellar cortex of an L7-YC2.60 mouse (#1055). (G) A bright field image of the dorsal surface of the cerebellum of a living L7-YC2.60 mouse (#6709). Scale bar: 1 mm. (H) Venus fluorescence image of the dorsal surface of the cerebellum of L7-YC2.60 mouse #1032. For comparison, a field-of-view (0.2025 mm^2^) of ordinary two-photon microscopy is shown by a red square. Scale bar: 1 mm. (I) Modified from Figure 1D. Each segment was transferred to 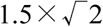 times outward from the center of the image while keeping its size. Because they are well separated, the individual segments are clearly delineated. A white broken line indicates the midline. Lobule names are abbreviated as in Figure 1B. Scale bar: 1 mm for the segment size.

**Figure S3.**
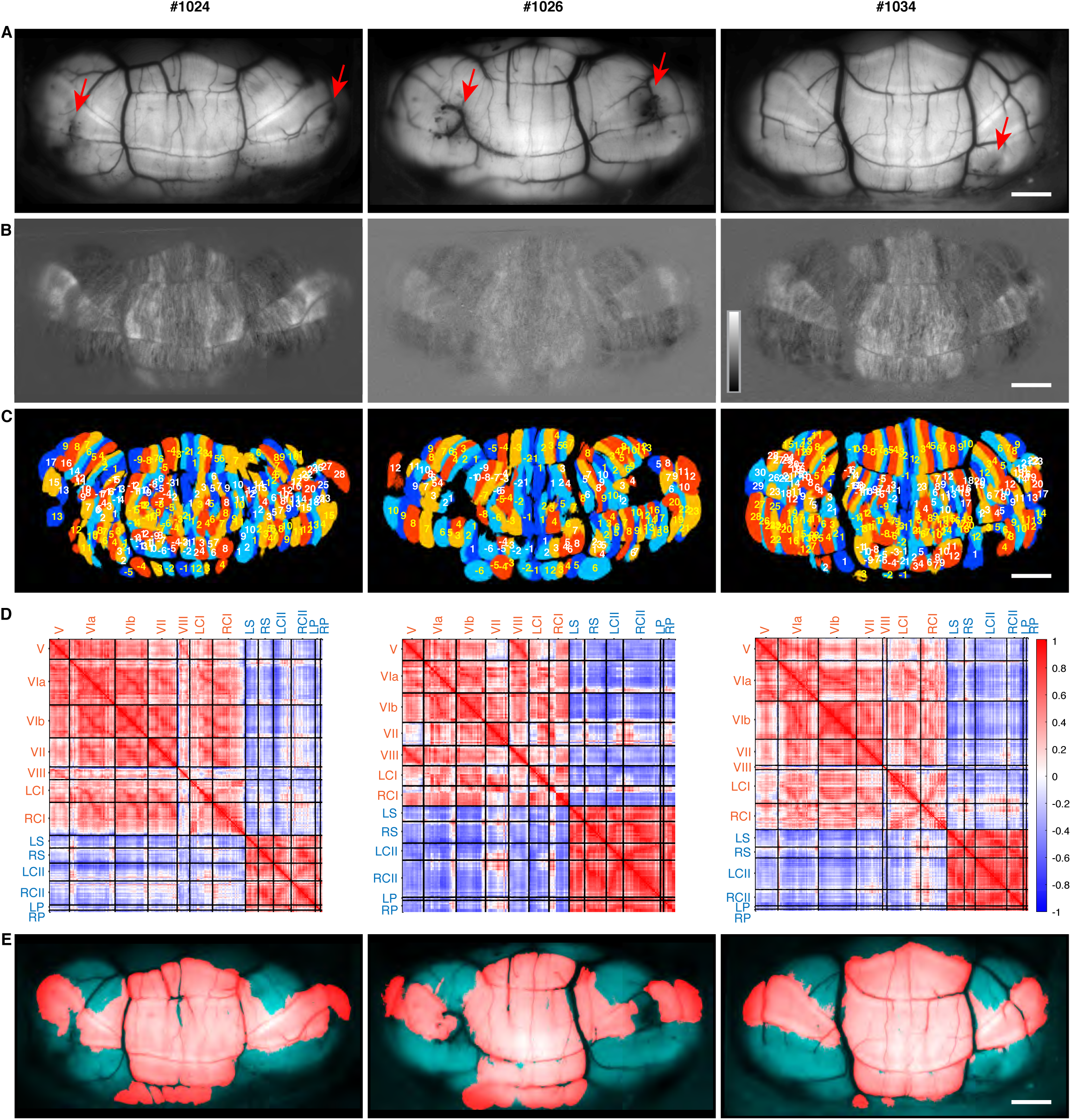
Olivocerebellar segments and cross characterized in three L7-YC2.60 mice (#1024, #1026, and #1034) (A) Venus fluorescence images indicating the existence of perioperative bleeding (red arrows). Refer to Figure 1A for #1032. (B) Snapshots of mean-subtracted ratio images (16-bit). Consecutive three frames were averaged. Refer to Figure 1C for #1032. Gray scale: 15,000–42,000 for #1024, 17,000–26,000 for #1026 and 15,000–34,000 for #1034. (C) Olivocerebellar segments identified with the independent component analysis of spontaneous CS activity; a total of 203, 164, and 238 segments were identified for #1024, #1026 and #1034, respectively. They are colored, labeled, and numbered as for #1032 (Figure 1D). Segments corresponding to the bleeding sites (indicated in (A)) were almost missing. (D) Cross-correlation coefficients (time lag = 0) between segmental FRET signals, revealing the same lobule categorization as made for #1032 (Figure 2B). Thus, the olivocerebellar cross was detected reproducibly in the cerebella of mice #1024, #1026, and #1034. Lobule names are abbreviated and labeled as in Figure 2B. (E) Segments inside the olivocerebellar cross are highlighted in red. Refer to Figure 2E for #1032. Scale bars: 1 mm.

**Figure S4.**
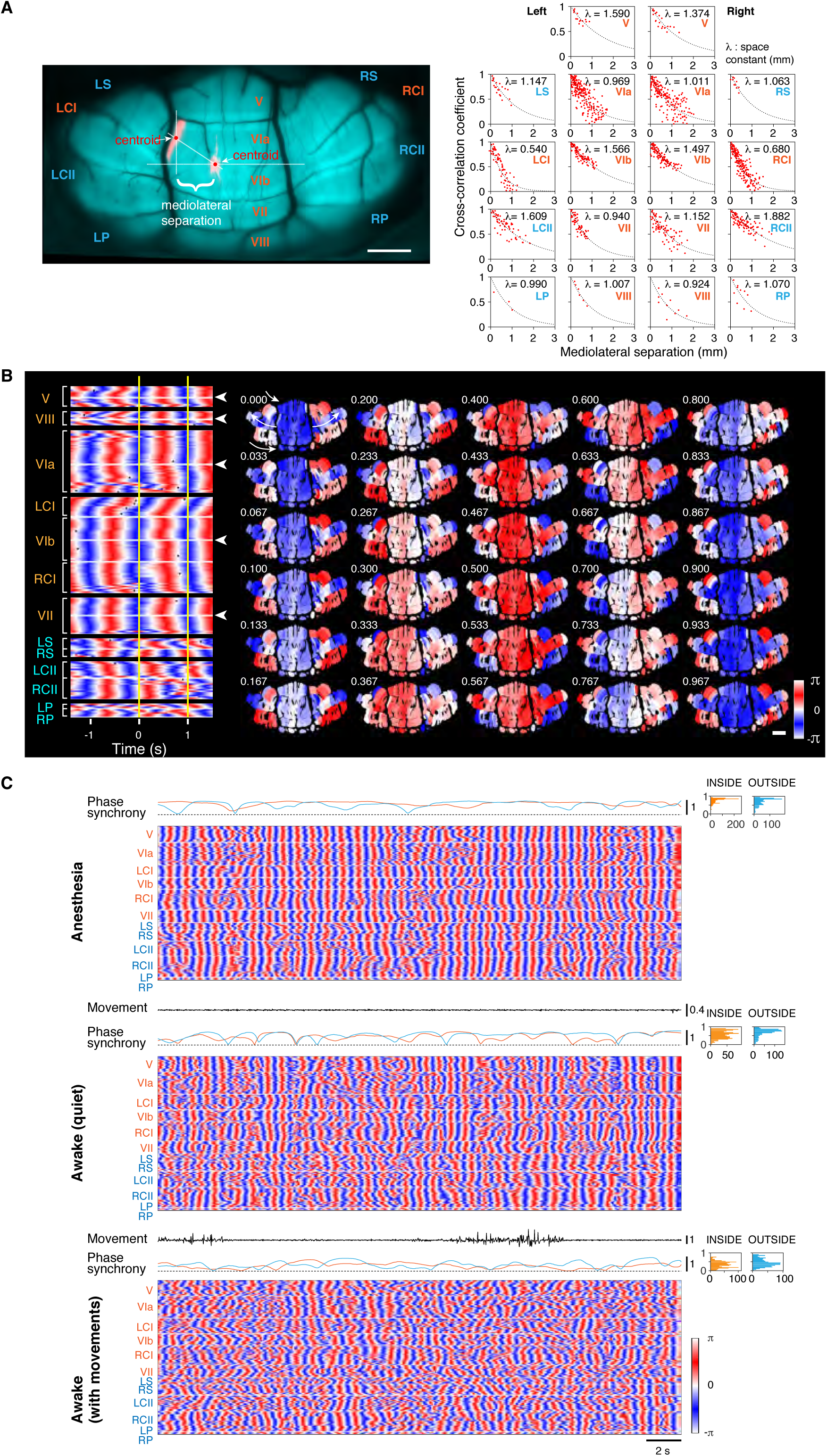
Local synchrony of CS occurrence and global dynamics of CS phase transition. (A) *left*, A fluorescence image of the dorsal surface of the cerebellum of L7-YC2.60 mouse #1032 with the names of examined lobules. The mediolateral separation distance between two olivocerebellar segments within VIa is shown as an example. Scale bar: 1 mm. *right*, Cross-correlation coefficients of spontaneous CS activity and mediolateral separation distances between two segments in each of the examined lobules are plotted. Each dot indicates an averaged cross-correlation coefficient from 25 measurements of 1-min recording. Dotted lines indicate an exponential function,

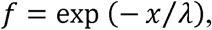

where λ is the space constant (mm) and *x* is the mediolateral separation distance (mm) between paired segments. See Table S2 for calculated λ values. The abbreviated names of lobules inside and outside the olivocerebellar cross are labelled with orange and cyan, respectively, as in Figure 2B. (B) *left*, The phase of spontaneous oscillations in CS activity of individual olivocerebellar segments was calculated (see Figure 2C) and is displayed with a circular color code. White arrowheads indicate the midlines of vermal lobules. *right*, Sequential images during a time period between two vertical yellow lines in the left panel. White arrows indicate directional flows of phase transitions, which moved outwardly inside the olivocerebellar cross but bi-directionally outside. Data were obtained from L7-YC2.60 mouse #1032. The oscillation frequencies statistically obtained from 25 measurements with this mouse were 1.15 ± 0.02 Hz (151 IN segments) and 1.12 ± 0.03 Hz (45 OUT segments). Likewise, the frequency values were obtained from three other mice; 1.22 ± 0.03 Hz (148 IN segments) and 1.11 ± 0.05 Hz (55 OUT segments) were calculated from 26 measurements with #1024; 1.03 ± 0.01 Hz (101 IN segments) and 1.02 ± 0.02 Hz (63 OUT segments) were calculated from 19 measurements with #1026; and 1.11 ± 0.04 Hz (169 IN segments) and 1.06 ± 0.05 Hz (69 OUT segments) were calculated from 12 measurements with #1034. On the whole, the spontaneous oscillations at around 1 Hz were found to occur uniformly in the cerebellar cortex of all the analyzed mice. (C) State dependent changes of the phase synchrony of segmental CS activities over the cerebellar cortex. Data were obtained from an L7-YC2.60 mouse (#6702) under three different conditions: Anesthesia, Awake (quiet), and Awake (with movements). The phase of spontaneous oscillations was calculated for individual segments as in Figure 2C and is displayed with a circular color code. The degree of the phase synchrony (see Methods) among segments inside (orange line) or outside (cyan line) the olivocerebellar cross is shown as a temporal profile (left) and a histogram (right). The movement of the mouse was measured by taking the first derivative of the average intensity of the face and forelimbs in an infrared movie. (B and C) Segments are aligned as in Figure 2A. Lobule names are abbreviated and labeled as in Figure 2B.

**Figure S5.**
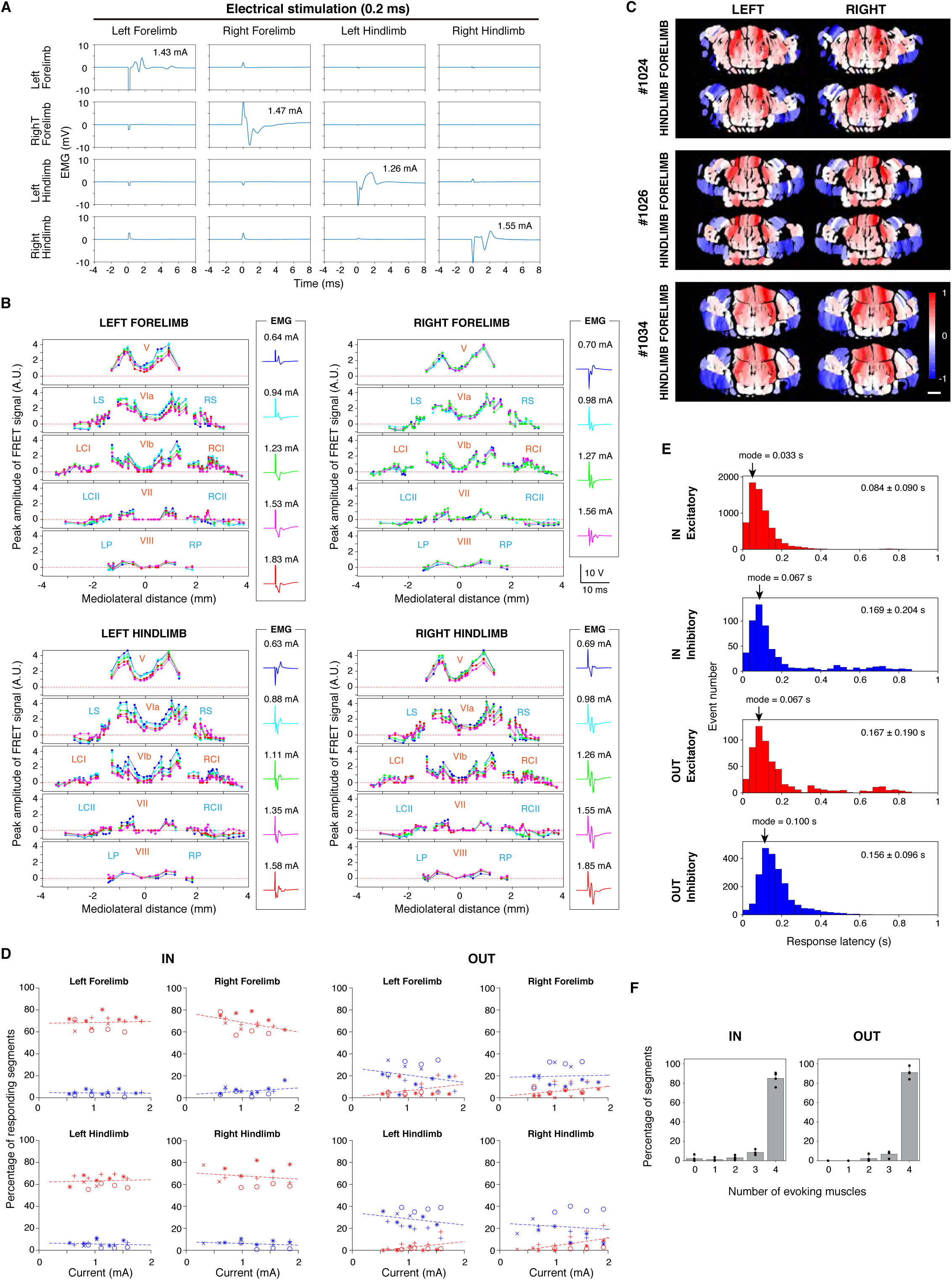
CS responses evoked by limb muscle stimulations. (A) Electromyographical examination of the specificity of muscle stimulation. Stimulation of each of the four limb muscles was repeated 30 times. Averaged traces that were recorded from all the four limb muscles are shown. These data were obtained from the experiments shown in Figure 3C. (B) Intensity dependency of evoked CS responses. Peak amplitudes of averaged CS responses (n = 30) evoked by several stimuli of different intensities are plotted as a function of mediolateral distance. Data were obtained from the L7-YC2.60 mouse (#1032). Lobule names are abbreviated and labeled as in Figure 2B. (C) Patterns of evoked CS responses observed in #1024, #1026, and #1034 were similar to those in Figure 3C. #1024: Peak amplitudes of averaged CS signals (n = 30) were mapped for the stimulation of the biceps brachii of the left forelimb (1.53 mA) (top left), biceps brachii of the right forelimb (1.27 mA) (top right), biceps femoris of the left hindlimb (1.35 mA) (bottom left) and biceps femoris of the right hindlimb (1.55 mA) (bottom right). #1026: Peak amplitudes of averaged CS signals (n = 30) were mapped for the stimulation of the biceps brachii of the left forelimb (1.23 mA) (top left), biceps brachii of the right forelimb (1.17 mA) (top right), biceps femoris of the left hindlimb (1.35 mA) (bottom left) and biceps femoris of the right hindlimb (1.55 mA) (bottom right). #1034: Peak amplitudes of averaged CS signals (n = 30) were mapped for the stimulation of the biceps brachii of the left forelimb (0.94 mA) (top left), biceps brachii of the right forelimb (0.98 mA) (top right), biceps femoris of the left hindlimb (1.03 mA) (bottom left) and biceps femoris of the right hindlimb (0.59 mA) (bottom right). Positive/negative responses (*i*.*e*. increase/decrease in the number of CS-generating PCs within a segment) were normalized against the maximum/minimum values and are colored in red/blue, respectively. Scale bar: 1 mm. See Figure 3C for the results from L7-YC2.60 mouse #1032. (D) Percentages of segments showing excitatory (increase in CS activity, red) or inhibitory (decrease in CS activity, blue) responses are plotted against stimulation intensities. Data from four different L7-YC2.60 mice (#1024, #1026, #1032, and #1034) are shown with different symbols. Pearson’s r values (P values) are as follows. IN: −0.037 (0.887), 0.313 (0.237), −0.108 (0.692) and −0.269 (0.314) for excitatory responses to the left forelimb, right forelimb, left hindlimb and right hindlimb, respectively, and 0.061 (0.817), −0.452 (0.079), 0.071 (0.793) and −0.185 (0.493) for inhibitory responses to the left forelimb, right forelimb, left hindlimb and right hindlimb, respectively. OUT: −0.361 (0.154), 0.338 (0.201), 0.421 (0.105) and 0.535 (0.033) for excitatory responses to the left forelimb, right forelimb, left hindlimb and right hindlimb, respectively, and −0.255 (0.323), 0.041 (0.881), −0.225 (0.402) and −0.123 (0.651) for inhibitory responses to the left forelimb, right forelimb, left hindlimb and right hindlimb, respectively. A low P value (< 0.05, high statistical significance) was obtained for the linear regression for excitatory responses outside the olivocerebellar cross to right hindlimb stimulation. Degrees of freedom equal 15. (E) Histograms of response latency of excitatory and inhibitory CS responses observed in IN and OUT segments after muscle stimulation. Response latency was determined as the minimum time at which the distributions of evoked and basal FRET signals were distinguished statistically by the Kolmogorov-Smirnov test. Mean ± standard deviation is shown in each histogram, calculated from 2,400 trials for all the four limb muscles of four L7-YC2.60 mice (#1024, #1026, #1032 and #1034). (F) Percentages of olivocerebellar segments (IN and OUT) receiving inputs from 0, 1 2, 3, and 4 limb muscles. The existence of input in each segment was determined by comparing two distributions of the peak amplitude between evoked and basal FRET signals. The Kolmogorov-Smirnov test was used to quantify the equality of two distributions. Mean values obtained from four L7-YC2.60 mice (#1024, #1026, #1032 and #1034) are shown (total 2,340 trials). Error bars correspond to the standard deviation. (D–F) IN: segments inside the olivocerebellar cross. OUT: segments outside the olivocerebellar cross.

**Figure S6.**
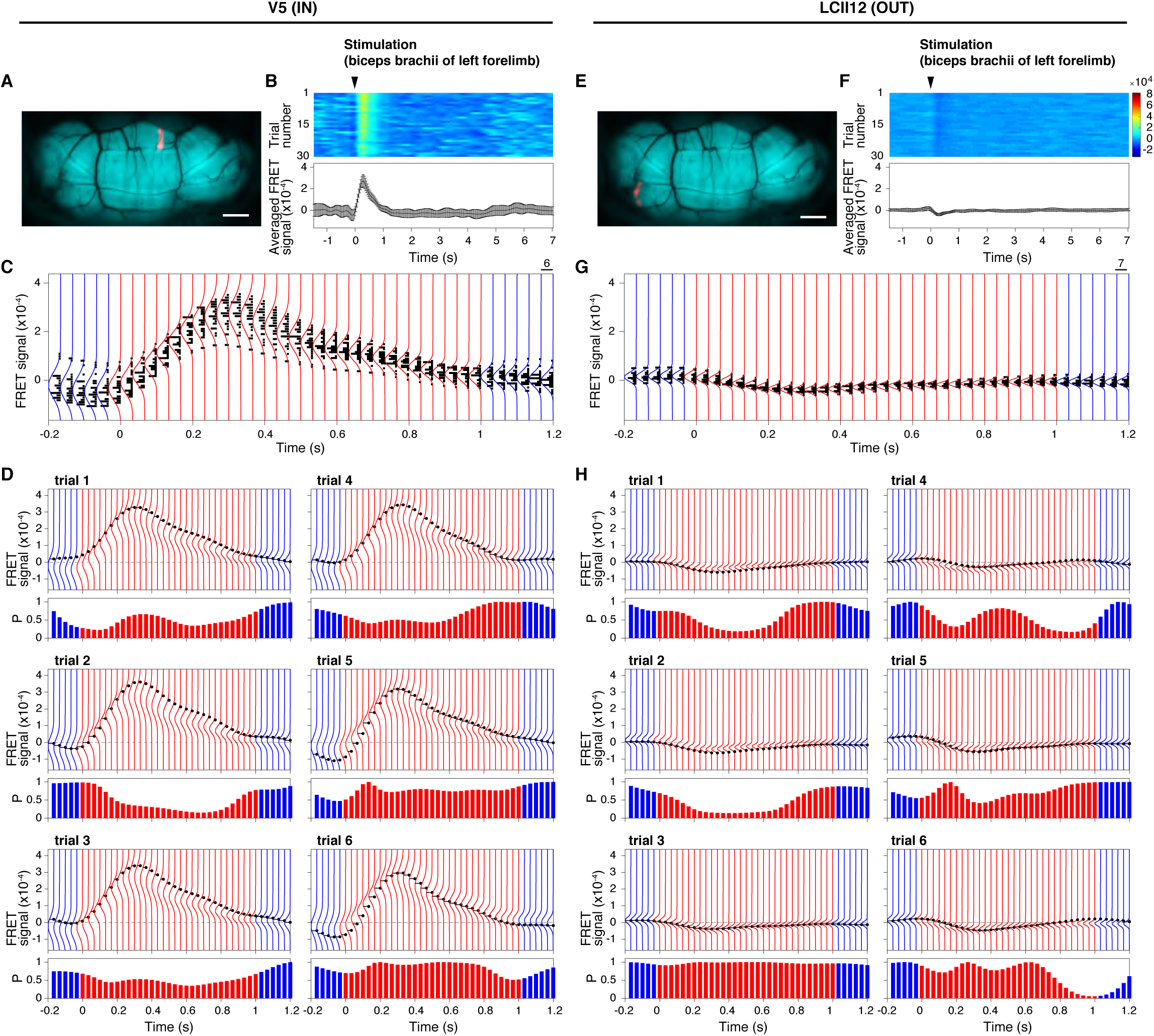
Conditional probabilities of FRET signals observed in single segments with multiple trials of muscle stimulation (left forelimb) (A–D) V5 is an olivocerebellar segment located inside the olivocerebellar cross. (E–H) LCII12 is an olivocerebellar segment located outside the olivocerebellar cross. (A and E) The segments are highlighted in magenta. Scale bars: 1 mm. (B and F) Evoked FRET signals during 30 trials of stimulation of biceps brachii of left forelimb. The stimulus was given at time 0 (duration: 0.2 msec, intensity: 1.26 mA). Time courses of color-coded FRET signals are shown individually (top). Average time courses are shown with error bars that represent standard deviations (bottom). (C and G) Histograms (horizontal black bars) of FRET signals observed at various time points. Probability density functions were estimated by fitting a Gaussian distribution function to each histogram (red lines in the time range from 0 to 1 sec and blue lines for others). (D and H) FRET signals (black dots) observed in the first six consecutive trials (B and F) and the probability density functions (C and G) are superimposed. Conditional probability (P) of FRET signals, calculated in comparison with the probability density functions (horizontal black bars), are shown by vertical bars (red, between 0 and 1 sec; blue, before 0 sec and after 1 sec). The conditional probability was normalized against the maximum value. Data were obtained from L7-YC2.60 mouse #1032.

**Figure S7.**
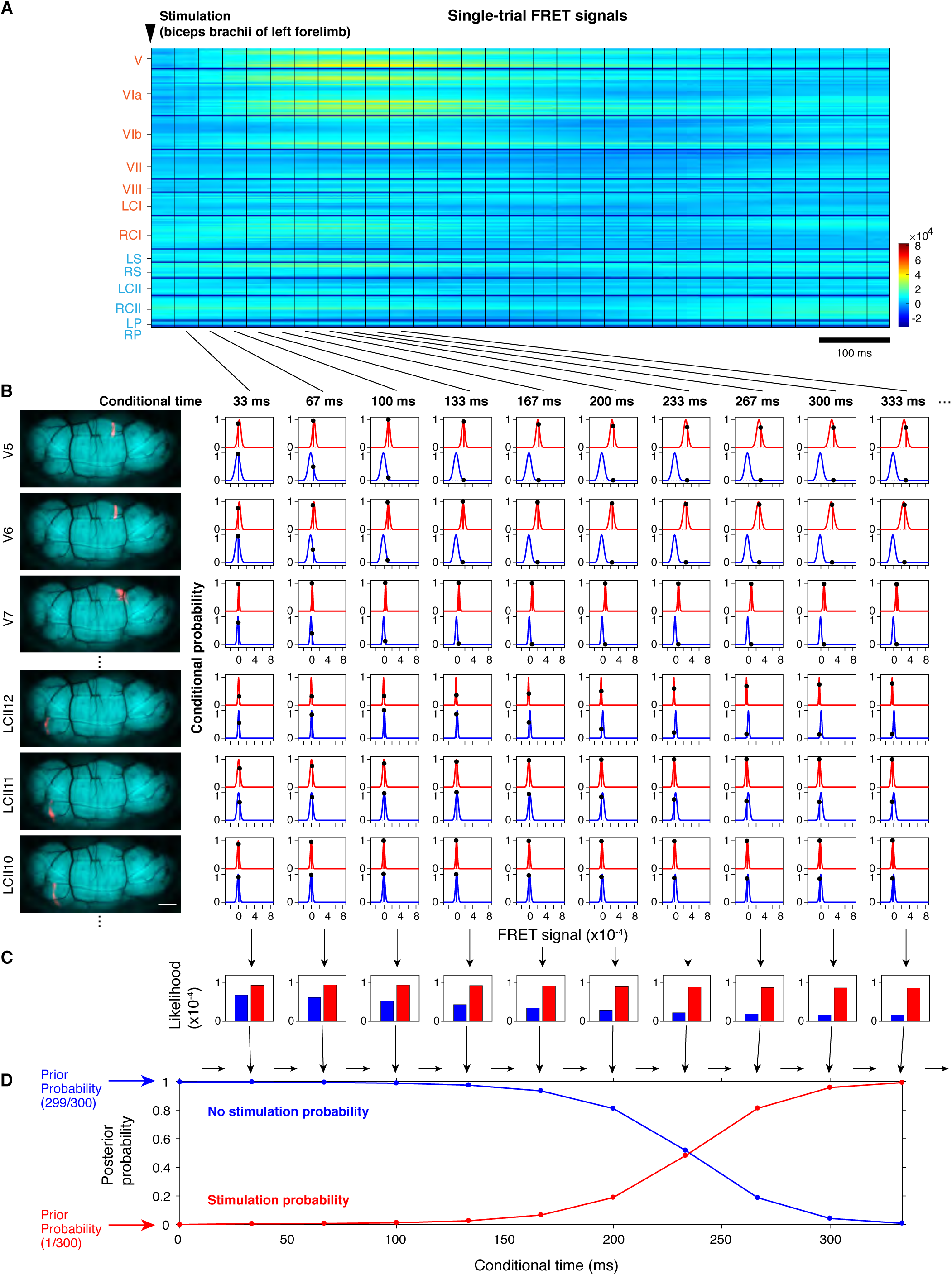
Cerebellum-wide calculation of posterior probabilities (stimulation probabilities) using the Bayesian updating algorithm. (A) Time courses of color-coded FRET signals during a single trial observed in all segments. At conditional time = 0 (indicated by a downward arrowhead), an electrical stimulus (duration: 0.2 msec, intensity: 1.83 mA) was applied to the biceps brachii of left forelimb. Abbreviated names of lobules inside and outside the olivocerebellar cross are labeled with orange and cyan, respectively as in Figure 2B. The same experiment as shown in Figure 4, but showing another trial. (B) Conditional probabilities of observed FRET signals (black dots) given the occurrence and nonoccurrence of stimulation were measured in each segment at each conditional time point from the estimated probability density functions of evoked (red) and basal (blue) FRET signals of multi-trial experiments, respectively (see Figure S6). Data from three segments inside the olivocerebellar cross (V5, V6, and V7) and three segments outside (LCII10, LCII11, and LCII12) are shown as examples. The six segments are individually highlighted in magenta. Scale bar: 1 mm. (C) Likelihoods for conditions of the occurrence (red) and nonoccurrence (blue) at each conditional time point. They were calculated by multiplying the conditional probabilities for all the segments. (D) Time-dependent posterior probabilities of the occurrence (red) and nonoccurrence (blue) of the stimulation obtained by the recursive calculation. At conditional time = 0, 1/300 and 299/300 were adopted as the prior probabilities of occurrence and nonoccurrence, respectively; each experiment had 300 measurements. Data were obtained from L7-YC2.60 mouse #1024.

**Figure S8.**
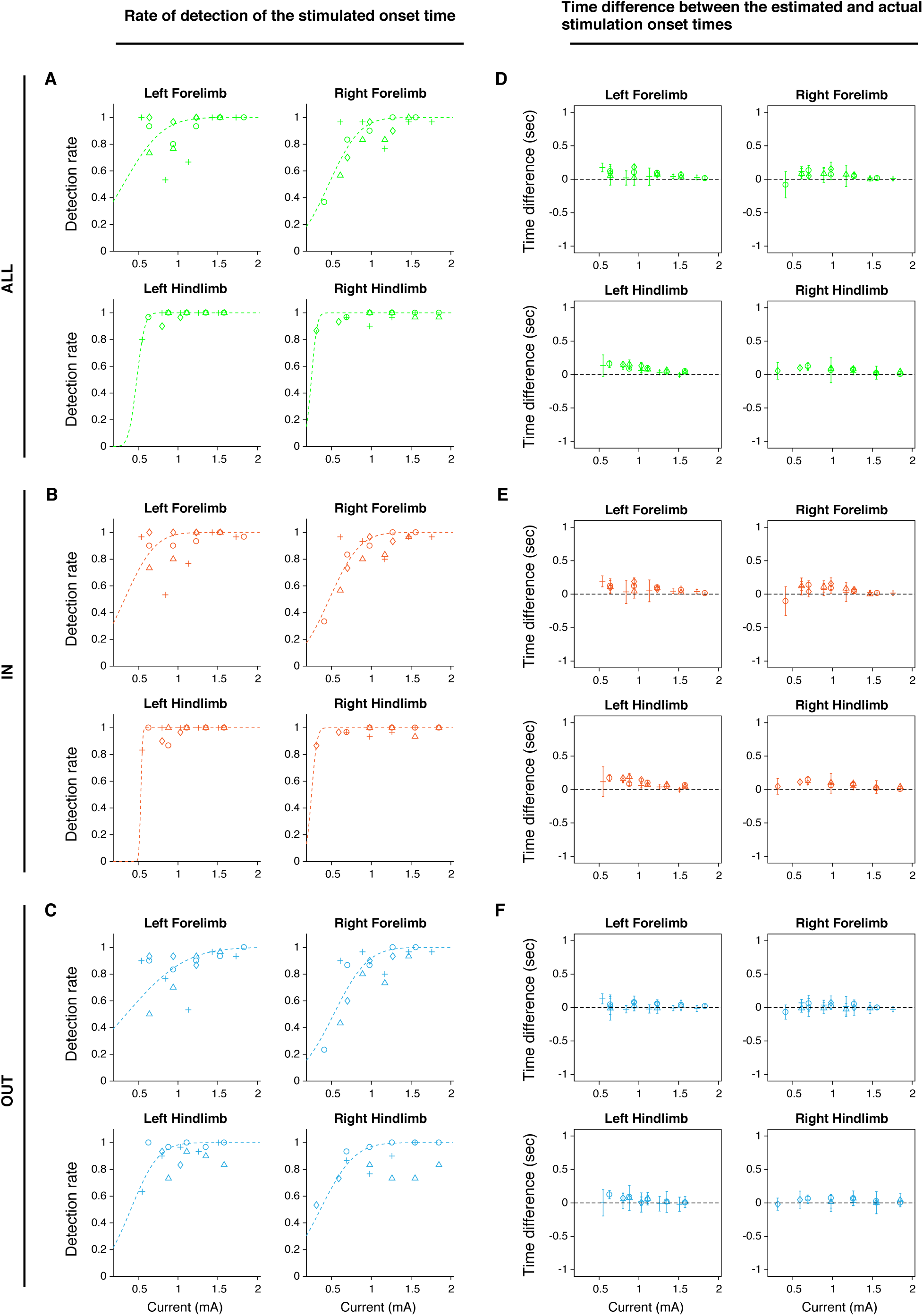
Efficiency and accuracy of estimation of the stimulation onset time. (A–C) Detection rate vs. current intensity. Relationships between the current applied to the stimulated muscle and the rate of detection of the stimulation onset time for all segments (A), segments inside the olivocerebellar cross (B), and segments outside the cross (C). Each data point is the rate of detected trials out of 30. The broken line indicates a normal cumulative distribution function, *p*,

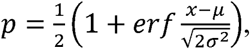

where *μ* is the mean and *σ*^2^ is the variance, fit to the detection rate *x*. (D–F) Time difference between the estimated and actual stimulation onset time for all segments (D), segments inside the olivocerebellar cross (E) and segments outside the cross (F). Each data point is a mean ± standard deviation (n = 30). Statistical values for all four limb muscles (mean ± standard deviation; n = 1,920 from four mice) are 0.0647 ± 0.0887 sec for all segments, 0.0736 ± 0.0903 sec for segments inside the olivocerebellar cross, and 0.0156 ± 0.1218 sec for segments outside the cross. (A–F) Data obtained from four L7-YC2.60 mice (#1024, #1026, #1032 and #1034) are shown as circles, triangles, crosses, and diamonds, respectively.

**Table S1.** Animals used in this study.

| Mouse | Strain | Calcium sensor | Age | Figure | Table |
| --- | --- | --- | --- | --- | --- |
| #127 | ICR | YC2.60<sup>*1</sup> | P186 | S2A, S2B, S2C, S2D |  |
| #1024 | C57Black6J | L7-Cre::tLSL-YC2.60<sup>*2</sup> | P96 | 4, 5, S3, S5C, S5D, S5E,<br>S5F, S6, S7, S8 | 1, S2 |
| #1026 | C57Black6J | L7-Cre::tLSL-YC2.60<sup>*2</sup> | P96 | 4C, 5B, 5C, 5D, S3, S5C,<br>S5D, S5E, S5F, S8 | 1, S2 |
| #1032 | C57Black6J | L7-Cre::tLSL-YC2.60<sup>*2</sup> | P84 | 1, 2, 3, 4C, S2H, S2I, S4A,<br>S4B, S5A, S5B, S5D,<br>S5E, S5F, S8 | S2 |
| #1034 | C57Black6J | L7-Cre::tLSL-YC2.60<sup>*2</sup> | P88 | 4C, 5B, 5C, 5D, S3, S5C,<br>S5D, S5E, S5F, S8 | 1, S2 |
| #1055 | C57Black6J | L7-Cre::tLSL-YC2.60<sup>*2</sup> | P182 | S2F |  |
| #6702 | C57Black6J | L7-Cre::tLSL-YC2.60<sup>*2</sup> | P58, 59 | S4C | S2 |
| #6709 | C57Black6J | L7-Cre::tLSL-YC2.60<sup>*2</sup> | P61 | S2G |  |

<sup>\*1</sup>: *in utero* injection of recombinant adenovirus vector
<sup>\*2</sup>: transgenic mouse line

**Table S2.**
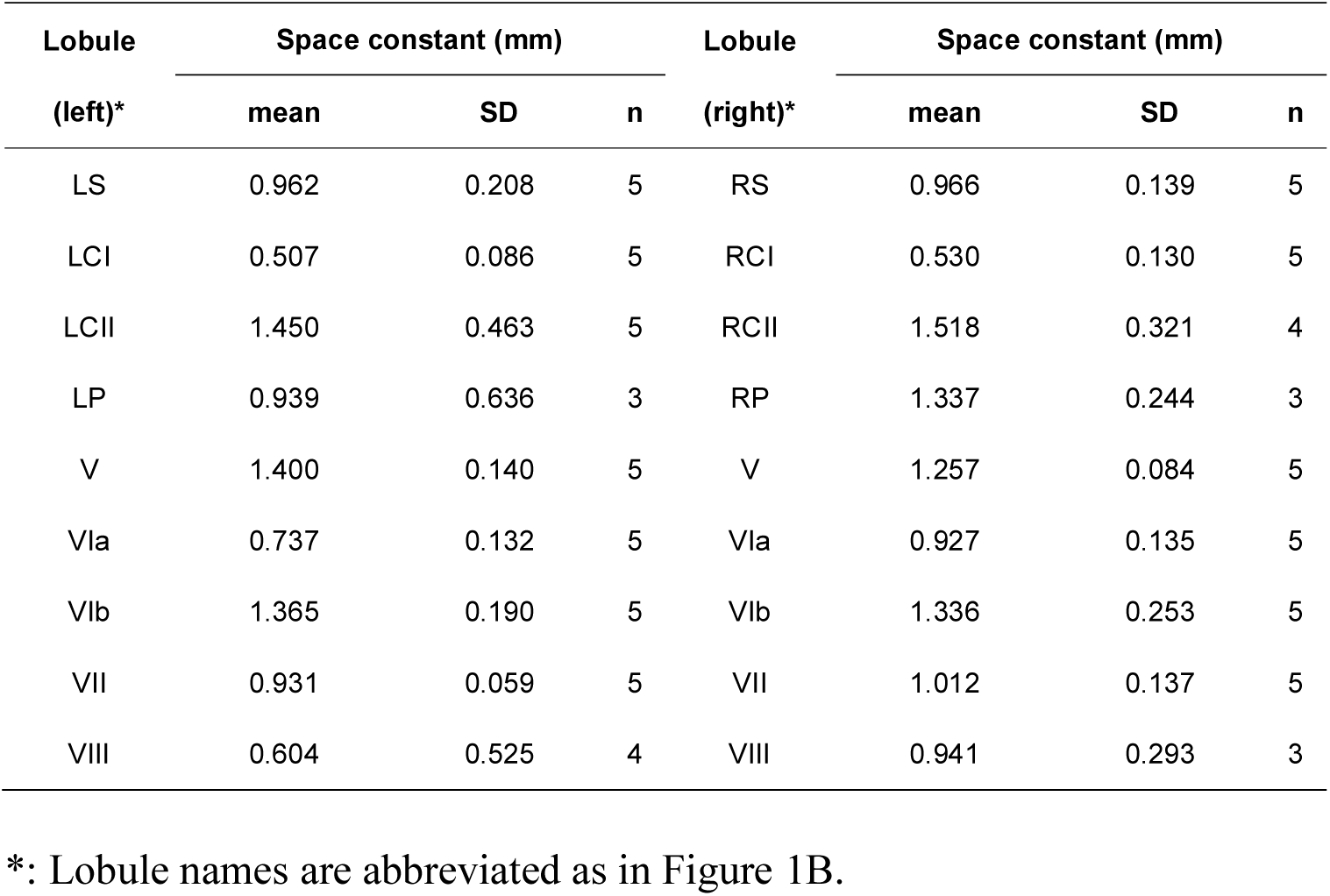
Space constant for CS synchrony.

**Table S3.**
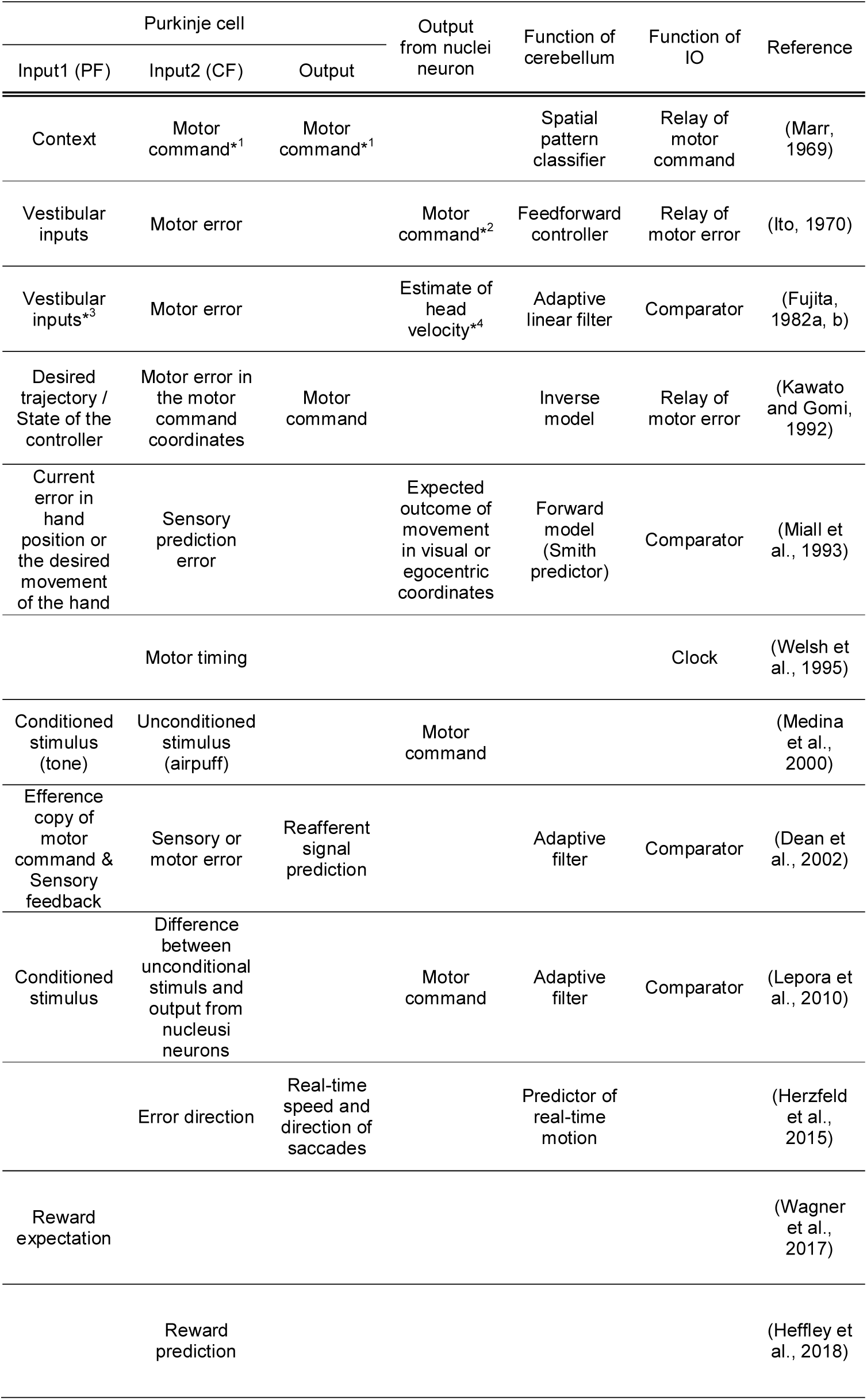
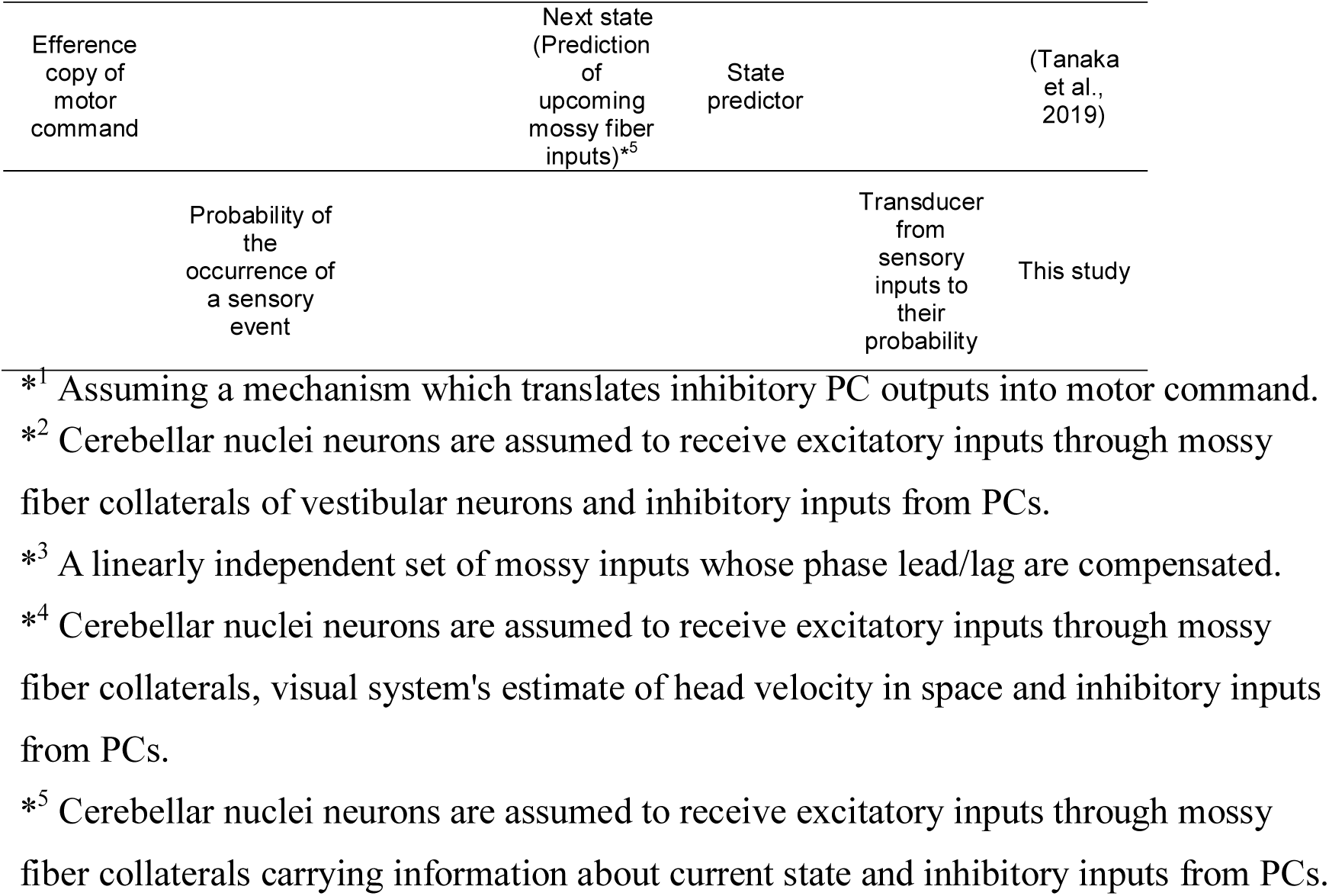
Proposals for the input and output relationship of the cerebellum.

## Notes

### Competing Interest Statement

The authors have declared no competing interest.

## REFERENCES

Andersson, G., and Oscarsson, O. (1978). Climbing fiber microzones in cerebellar vermis and their projection to different groups of cells in the lateral vestibular nucleus. Exp Brain Res 32, 565–579.

Apps, R., and Garwicz, M. (2005). Anatomical and physiological foundations of cerebellar information processing. Nat Rev Neurosci 6, 297–311.

Apps, R., and Hawkes, R. (2009). Cerebellar cortical organization: a one-map hypothesis. Nat Rev Neurosci 10, 670–681.

Baumann, O., Borra, R.J., Bower, J.M., Cullen, K.E., Habas, C., Ivry, R.B., Leggio, M., Mattingley, J.B., Molinari, M., Moulton, E.A., et al. (2015). Consensus paper: the role of the cerebellum in perceptual processes. Cerebellum 14, 197–220.

Catz, N., Dicke, P.W., and Thier, P. (2005). Cerebellar complex spike firing is suitable to induce as well as to stabilize motor learning. Curr Biol 15, 2179–2189.

Combs, C.M. (1954). Electro-anatomical study of cerebellar localization; stimulation of various afferents. J Neurophysiol 17, 123–143.

Dean, P., Porrill, J., Ekerot, C.F., and Jörntell, H. (2010). The cerebellar microcircuit as an adaptive filter: experimental and computational evidence. Nat Rev Neurosci 11, 30–43.

Eccles, J.C., Ito, M., and Szentágothai, J. (1967). The Cerebellum as a Neuronal Machine (Berlin, Heidelberg: Springer).

Eccles, J.C., Provini, L., Strata, P., and Taborikova, H. (1968). Topographical investigations on the climbing fiber inputs from forelimb and hindlimb afferents to the cerebellar anterior lobe. Exp Brain Res 6, 195–215.

Ekerot, C.F., and Larson, B. (1979). The dorsal spino-olivocerebellar system in the cat. II. Somatotopical organization. Exp Brain Res 36, 219–232.

Friston, K. (2005). A theory of cortical responses. Philos Trans R Soc Lond B Biol Sci 360, 815–836.

Friston, K. (2010). The free-energy principle: a unified brain theory? Nat Rev Neurosci 11, 127–138.

Horikawa, K., Yamada, Y., Matsuda, T., Kobayashi, K., Hashimoto, M., Matsu-ura, T., Miyawaki, A., Michikawa, T., Mikoshiba, K., and Nagai, T. (2010). Spontaneous network activity visualized by ultrasensitive Ca^2+^ indicators, yellow Cameleon-Nano. Nat Methods 7, 729–732.

Ito, M. (1984). The Cerebellum and Neural Control (Raven Press).

Kandel, E., Schwartz, J., Jessell, T., Siegelbaum, S., and Hudspeth, A.J. (2012). Principles of Neural Science (McGraw-Hill Professional).

Keating, J.G., and Thach, W.T. (1995). Nonclock behavior of inferior olive neurons: interspike interval of Purkinje cell complex spike discharge in the awake behaving monkey is random. J Neurophysiol 73, 1329–1340.

Kitazawa, S., Kimura, T., and Yin, P.B. (1998). Cerebellar complex spikes encode both destinations and errors in arm movements. Nature 392, 494–497.

Knill, D.C., and Pouget, A. (2004). The Bayesian brain: the role of uncertainty in neural coding and computation. Trends Neurosci 27, 712–719.

Kuroki, S., Yoshida, T., Tsutsui, H., Iwama, M., Ando, R., Michikawa, T., Miyawaki, A., Ohshima, T., and Itohara, S. (2018). Excitatory neuronal hubs configure multisensory integration of slow waves in association cortex. Cell Rep 22, 2873–2885.

Lang, E.J., Apps, R., Bengtsson, F., Cerminara, N.L., De Zeeuw, C.I., Ebner, T.J., Heck, D.H., Jaeger, D., Jörntell, H., Kawato, M., et al. (2017). The Roles of the Olivocerebellar Pathway in Motor Learning and Motor Control. A Consensus Paper. Cerebellum 16, 230–252.

Leicht, R., and Schmidt, R.F. (1977). Somatotopic studies on the vermal cortex of the cerebellar anterior lobe of unanaesthetized cats. Exp Brain Res 27, 479–490.

Llinás, R., and Sasaki, K. (1989). The Functional Organization of the Olivo-Cerebellar System as Examined by Multiple Purkinje Cell Recordings. Eur J Neurosci 1, 587–602.

Makino, H., Ren, C., Liu, H., Kim, A.N., Kondapaneni, N., Liu, X., Kuzum, D., and Komiyama, T. (2017). Transformation of Cortex-wide Emergent Properties during Motor Learning. Neuron 94, 880–890.

Manni, E., and Petrosini, L. (2004). A century of cerebellar somatotopy: a debated representation. Nat Rev Neurosci 5, 241–249.

Marr, D. (1969). A theory of cerebellar cortex. J Physiol 202, 437–470.

Miyawaki, A., Llopis, J., Heim, R., McCaffery, J.M., Adams, J.A., Ikura, M., and Tsien, R.Y. (1997). Fluorescent indicators for Ca^2+^ based on green fluorescent proteins and calmodulin. Nature 388, 882–887.

Mukamel, E.A., Nimmerjahn, A., and Schnitzer, M.J. (2009). Automated analysis of cellular signals from large-scale calcium imaging data. Neuron 63, 747–760.

Nagai, T., Yamada, S., Tominaga, T., Ichikawa, M., and Miyawaki, A. (2004). Expanded dynamic range of fluorescent indicators for Ca^2+^ by circularly permuted yellow fluorescent proteins. Proc Natl Acad Sci U S A 101, 10554–10559.

Oscarsson, O. (1968). Termination and functional organization of the ventral spino-olivocerebellar path. J Physiol 196, 453–478.

Oscarsson, O. (1980). Functional organization of olivary projection to the cerebellar anterior lobe. In The Inferior Olivary Nucleus, Anatomy and Physiology, C.d.M. Jacques Courville, Yves Lamarre, ed. (New York: Raven Press), pp. 279–289.

Penfield, W., and Rasmussen, T. (1950). The Cerebral Cortex of Man (New York: MacMillan).

Purves, D., Augustine, G.J., Fitzpatrick, D., Hall, W.C., LaMantia, A.-S., Mooney, R.D., Platt, M.L., and White, L.E. (2018). Neuroscience, 6th ed. (Sinauer Associates).

Purves, D., Brannon, E.M., Cabeza, R., Huettel, S.A., and LaBar, K.S. (2007). Principles of Cognitive Neuroscience (Sinauer Associates).

Rao, R.P., and Ballard, D.H. (1999). Predictive coding in the visual cortex: a functional interpretation of some extra-classical receptive-field effects. Nat Neurosci 2, 79–87.

Robertson, L.T., Laxer, K.D., and Rushmer, D.S. (1982). Organization of climbing fiber input from mechanoreceptors to lobule V vermal cortex of the cat. Exp Brain Res 46, 281–291.

Rondi-Reig, L., Paradis, A.L., Lefort, J.M., Babayan, B.M., and Tobin, C. (2014). How the cerebellum may monitor sensory information for spatial representation. Front Syst Neurosci 8, 205.

Shadmehr, R., Smith, M.A., and Krakauer, J.W. (2010). Error correction, sensory prediction, and adaptation in motor control. Annu Rev Neurosci 33, 89–108.

Snider, R.S., and Stowell, A. (1944). Receiving areas of the tactile, auditory and visual systems in the cerebellum. J Neurophysiol 7, 331–358.

Von Helmholtz, H. (2005). Treatise on Physiological Optics, Vol. 3 (Courier Corporation).

Voogd, J., Shinoda, Y., Ruigrok, T.J., and Sugihara, I. (2013). Cerebellar nuclei and the inferior olivary nuclei: organization and connections. In Handbook of the Cerbellum and Cerebellar Disorders, M. Manto, D.L. Gruol, J.D. Schmahmann, N. Koibuchi, and F. Rossi, eds. (Springer), pp. 377–436.

Wolpert, D.M., Miall, R.C., and Kawato, M. (1998). Internal models in the cerebellum. Trends Cogn Sci 2, 338–347.

Woolsey, C.N., and Fairman, D. (1946). Contralateral, ipsilateral, and bilateral representation of cutaneous receptors in somatic areas I and II of the cerebral cortex of pig, sheep, and other mammals. Surgery 19, 684–702.

Yamada, Y., Michikawa, T., Hashimoto, M., Horikawa, K., Nagai, T., Miyawaki, A., Häusser, M., and Mikoshiba, K. (2011). Quantitative comparison of genetically encoded Ca indicators in cortical pyramidal cells and cerebellar Purkinje cells. Front Cell Neurosci 5, 18.

Zhang, K., Ginzburg, I., McNaughton, B.L., and Sejnowski, T.J. (1998). Interpreting neuronal population activity by reconstruction: unified framework with application to hippocampal place cells. J Neurophysiol 79, 1017–1044.

## References

Chaumont, J., Guyon, N., Valera, A.M., Dugue, G.P., Popa, D., Marcaggi, P., Gautheron, V., Reibel-Foisset, S., Dieudonne, S., Stephan, A., et al. (2013). Clusters of cerebellar Purkinje cells control their afferent climbing fiber discharge. Proc Natl Acad Sci U S A 110, 16223–16228.

Dean, P., Porrill, J., and Stone, J.V. (2002). Decorrelation control by the cerebellum achieves oculomotor plant compensation in simulated vestibulo-ocular reflex. Proc Biol Sci 269, 1895–1904.

Fujita, M. (1982a). Adaptive filter model of the cerebellum. Biol Cybern 45, 195–206.

Fujita, M. (1982b). Simulation of adaptive modification of the vestibulo-ocular reflex with an adaptive filter model of the cerebellum. Biol Cybern 45, 207–214.

Heffley, W., Song, E.Y., Xu, Z., Taylor, B.N., Hughes, M.A., McKinney, A., Joshua, M., and Hull, C. (2018). Coordinated cerebellar climbing fiber activity signals learned sensorimotor predictions. Nat Neurosci 21, 1431–1441.

Herzfeld, D.J., Kojima, Y., Soetedjo, R., and Shadmehr, R. (2015). Encoding of action by the Purkinje cells of the cerebellum. Nature 526, 439–442.

Ito, M. (1970). Neurophysiological aspects of the cerebellar motor control system. Int J Neurol 7, 162–176.

Kawato, M., and Gomi, H. (1992). A computational model of four regions of the cerebellum based on feedback-error learning. Biol Cybern 68, 95–103.

Lepora, N.F., Porrill, J., Yeo, C.H., and Dean, P. (2010). Sensory prediction or motor control? Application of marr-albus type models of cerebellar function to classical conditioning. Front Comput Neurosci 4, 140.

Medina, J.F., Garcia, K.S., Nores, W.L., Taylor, N.M., and Mauk, M.D. (2000). Timing mechanisms in the cerebellum: testing predictions of a large-scale computer simulation. J Neurosci 20, 5516–5525.

Miall, R.C., Weir, D.J., Wolpert, D.M., and Stein, J.F. (1993). Is the cerebellum a smith predictor? J Mot Behav 25, 203–216.

Mrsic-Flogel, T.D., Hofer, S.B., Ohki, K., Reid, R.C., Bonhoeffer, T., and Hubener, M. (2007). Homeostatic regulation of eye-specific responses in visual cortex during ocular dominance plasticity. Neuron 54, 961–972.

Ruigrok, T.J., and Voogd, J. (1990). Cerebellar nucleo-olivary projections in the rat: an anterograde tracing study with Phaseolus vulgaris-leucoagglutinin (PHA-L). J Comp Neurol 298, 315–333.

Snider, R.S. (1958). The cerebellum. Sci Am 199, 84–90.

Sugihara, I., Fujita, H., Na, J., Quy, P.N., Li, B.Y., and Ikeda, D. (2009). Projection of reconstructed single Purkinje cell axons in relation to the cortical and nuclear aldolase C compartments of the rat cerebellum. J Comp Neurol 512, 282–304.

Sugihara, I., and Shinoda, Y. (2004). Molecular, topographic, and functional organization of the cerebellar cortex: a study with combined aldolase C and olivocerebellar labeling. J Neurosci 24, 8771–8785.

Takita, K., Aoki, T., Sasaki, Y., Higuchi, T., and Kobayashi, K. (2003). High-accuracy subpixel image registration based on phase-only correlation. IEICE Transactions Fundamentals E86-A, 1925–1934.

Tanaka, H., Ishikawa, T., and Kakei, S. (2019). Neural Evidence of the Cerebellum as a State Predictor. Cerebellum 18, 349–371.

Uusisaari, M., and Knopfel, T. (2011). Functional classification of neurons in the mouse lateral cerebellar nuclei. Cerebellum 10, 637–646.

Wagner, M.J., Kim, T.H., Savall, J., Schnitzer, M.J., and Luo, L. (2017). Cerebellar granule cells encode the expectation of reward. Nature 544, 96–100.

Welsh, J.P., Lang, E.J., Suglhara, I., and Llinás, R. (1995). Dynamic organization of motor control within the olivocerebellar system. Nature 374, 453–457.

Yasuda, K., Hayashi, Y., Yoshida, T., Kashiwagi, M., Nakagawa, N., Michikawa, T., Tanaka, M., Ando, R., Huang, A., Hosoya, T., et al. (2017). Schizophrenia-like phenotypes in mice with NMDA receptor ablation in intralaminar thalamic nucleus cells and gene therapy-based reversal in adults. Transl Psychiatry 7, e1047.

